# Polybacterial Intracellular Macromolecules Shape Single-Cell Epikine Profiles in Upper Airway Mucosa

**DOI:** 10.1101/2024.10.08.617279

**Authors:** Quinn T. Easter, Zabdiel Alvarado-Martinez, Meik Kunz, Bruno Fernandes Matuck, Brittany T. Rupp, Theresa Weaver, Zhi Ren, Aleksandra Tata, Juan Caballero-Perez, Nick Oscarson, Akira Hasuike, Ameer N. Ghodke, Adam J. Kimple, Purushothama R. Tata, Scott H. Randell, Hyun Koo, Kang I. Ko, Kevin M. Byrd

## Abstract

The upper airway, particularly the nasal and oral mucosal epithelium, serves as a primary barrier for microbial interactions throughout life. Specialized niches like the anterior nares and the tooth are especially susceptible to dysbiosis and chronic inflammatory diseases. To investigate host-microbial interactions in mucosal epithelial cell types, we reanalyzed our single-cell RNA sequencing atlas of human oral mucosa, identifying polybacterial signatures (20% Gram-positive, 80% Gram-negative) within both epithelial- and stromal-resident cells. This analysis revealed unique responses of bacterial-associated epithelia when compared to two inflammatory disease states of mucosa. Single-cell RNA sequencing, *in situ* hybridization, and immunohistochemistry detected numerous persistent macromolecules from Gram-positive and Gram-negative bacteria within human oral keratinocytes (HOKs), including bacterial rRNA, mRNA and glycolipids. Epithelial cells with higher concentrations of *16S* rRNA and glycolipids exhibited enhanced receptor-ligand signaling *in vivo*. HOKs with a spectrum of polybacterial intracellular macromolecular (PIM) concentrations were challenged with purified exogenous lipopolysaccharide, resulting in the synergistic upregulation of select innate (*CXCL8*, *TNFSF15*) and adaptive (*CXCL17*, *CCL28*) epikines. Notably, endogenous lipoteichoic acid, rather than lipopolysaccharide, directly correlated with epikine expression *in vitro* and *in vivo*. Application of the Drug2Cell algorithm to health and inflammatory disease data suggested altered drug efficacy predictions based on PIM detection. Our findings demonstrate that PIMs persist within mucosal epithelial cells at variable concentrations, linearly driving single-cell effector cytokine expression and influencing drug responses, underscoring the importance of understanding host-microbe interactions and the implications of PIMs on cell behavior in health and disease at single-cell resolution.

**One-sentence summary:** This study reveals how persistent intracellular bacterial macromolecules in mucosal epithelial cells drive inflammatory signaling, offering new insights into microbial-host interactions and their potential impact on inflammatory disease treatment and drug efficacy.

## INTRODUCTION

Human beings are superorganisms that host numerous microbes. The upper airways, primarily comprising the nasal and oral cavities, are key interfaces between the internal and external environments and for microbial encounters; they are exposed daily to a variety of stressors, including environmental toxins, allergens, irritants, and complex microbial interactions*(1, 2)*. These interconnected tissues host a diverse microbiome of over 1,000 unique bacterial species, each colonizing biogeographical niches of the mucosa*(3–5)*. While most of these bacteria are Gram-positive, commensal, and not associated with disease, shifts in microbial communities— often involving both Gram-positive and Gram-negative species—can lead to dysbiosis*(6, 7)*. This imbalance sensitizes the host to chronic inflammatory conditions of the mucosa, such as chronic rhinosinusitis*(8)*, primary ciliary dyskinesia*(9)*, dental caries*(10)*, gingivitis, and periodontitis*(11)*. Understanding these mucosal-microbial dynamics at a single-cell and spatial resolution is critical for developing precision interventions for this wide range of dysbiosis-associated conditions*(12)*.

Further, distinct mineralized, mucosal, and biofluidic oral cavity niches support site-specific bacterial profiles*(13–15)*. A key specialized niche is the periodontium, which comprises alveolar bone, periodontal ligament, cementum, and gingival mucosa*(16)*. The periodontium is uniquely susceptible to chronic inflammation, termed either gingivitis or periodontitis*(17)*, both marked by significant gingival tissue inflammation which compromises the integrity of the mucosal epithelial barrier, secondary to bacterial dysbiosis in susceptible individuals. With over 80% of adults exhibiting signs of gingivitis, mucosal-microbial interactions are common and widespread. Left untreated, tissue breakdown progresses through immune and inflammatory responses, eventually reaching the alveolar bone*(18)*. Thus, early identification and intervention to modulate microbial influences on the mucosa are key to preventing local and systemic effects of chronic mucosal inflammatory diseases*(19)*.

Recently, we generated the first integrated single-cell RNA sequencing (scRNAseq) atlas of oral mucosa and periodontal ligament*(20)*. This atlas included samples from clinically healthy individuals and patients diagnosed with periodontitis*(21, 22)* or experimentally induced gingivitis*(23)*. As part of this study, we reannotated the scRNAseq data using unmapped reads, uncovering 37 unique bacterial RNA signals with cell-specific tropism. In structural cell types, such as epithelial and stromal cells, we identified cells containing mono- and polybacterial signatures—20% Gram-positive and 80% Gram-negative. Several Gram-negative periopathogens, including *Porphyromonas (P.) gingivalis, Tannerella forsythia, Treponema (T.) denticola,* and *Aggregatibacter actinomycetemcomitans*, have been extensively studied for their potential to invade epithelial cells in gingivitis and periodontitis*(24)*. Some Gram-positive bacteria like *Streptococcus sanguinis* and *Streptococcus mutans* are also implicated in periodontitis and exhibit some invasive capabilities*(25, 26)*. Moreover, components of Gram-positive bacteria, such as lipoteichoic acid (LTA), can induce inflammation, contributing to alveolar bone loss in periodontitis*(27)*. While our scRNAseq analysis detected polybacterial signals, we sought to determine whether intact Gram-positive and Gram-negative bacteria or their macromolecular components—RNA and glycolipids —were immunostimulatory to mucosal epithelial cells, regardless of disease state.

To address this, we further analyzed the scRNAseq atlas, focusing on primary human oral keratinocytes (HOKs) containing intracellular bacterial signatures. High-resolution confocal and transmission electron microscopy revealed minimal intact intracellular bacteria in HOKs from multiple donors. However, whole-genomic sequencing and multiplexed fluorescence *in situ* hybridization (FISH) with immunohistochemistry (IHC) identified numerous bacterial macromolecules within these cells, including LTA and lipopolysaccharides (LPS) over multiple passages. HOKs with higher concentrations of bacterial *16S* rRNA, LTA, and LPS exhibited enhanced receptor-ligand, or “epikine,” signaling. In conjunction with the CellChat algorithm, we observed that HOKs challenged with LPS synergistically upregulated specific innate immune epikines, such as *CXCL8* and *TNFSF15*, as well as adaptive immune epikines, like *CXCL17* and *CCL28*. Interestingly, LTA concentrations, rather than LPS, linearly correlated with innate epikine expression—both *in vitro* and *in vivo*. We then applied the Drug2Cell algorithm to predict drug efficacy based on polybacterial macromolecule detection*(28)*. The results suggested varying predicted drug interactions across 140 different compounds depending on the presence and concentration of these bacterial components in keratinocytes. Our findings demonstrate that polybacterial intracellular macromolecules (PIMs) can persist within oral mucosal epithelial cells at variable concentrations, driving single-cell effector cytokine expression and influencing predicted drug responses. This underscores the need to fully understand host-microbial interactions at the single-cell level, particularly within the upper airway mucosal tissues, to develop targeted therapies for mucosal inflammatory diseases with dysbiotic phenotypes.

## RESULTS

### Reanalysis of healthy and inflamed mucosal atlas reveals polybacterial signatures and their disease-agnostic impacts on host epithelium

The Human Cell Atlas Oral & Craniofacial Bionetwork efforts have already revealed heterogeneity among structural cell types of the oral and oropharyngeal tissues*(29)*. Related to this effort, we previously generated an integrated periodontal single-cell RNA sequencing (scRNAseq) atlas*(20)* (Figure 1a; publicly available at CZ CELLxGENE, ID: 71f4bccf-53d4-4c12-9e80-e73bfb89e398). This dataset 34-sample, 105918-cell atlas comprised a rich diversity of structural (i.e., epithelial-, stromal-resident) and immune (innate, adaptive) cell types. Furthermore, this atlas contained healthy, gingivitis, and periodontitis tissues. To ask questions about bacterial presence, diversity, and host impact associated with each cell type, we first modified, then applied the Single-cell Analysis of Host-Microbiome Interactions pipeline*(30)* (SAHMI; see: *Methods*) to the atlas, wherein we remapped scRNAseq reads that did not map to the human genome (Figure 1b).

**Figure 1.**
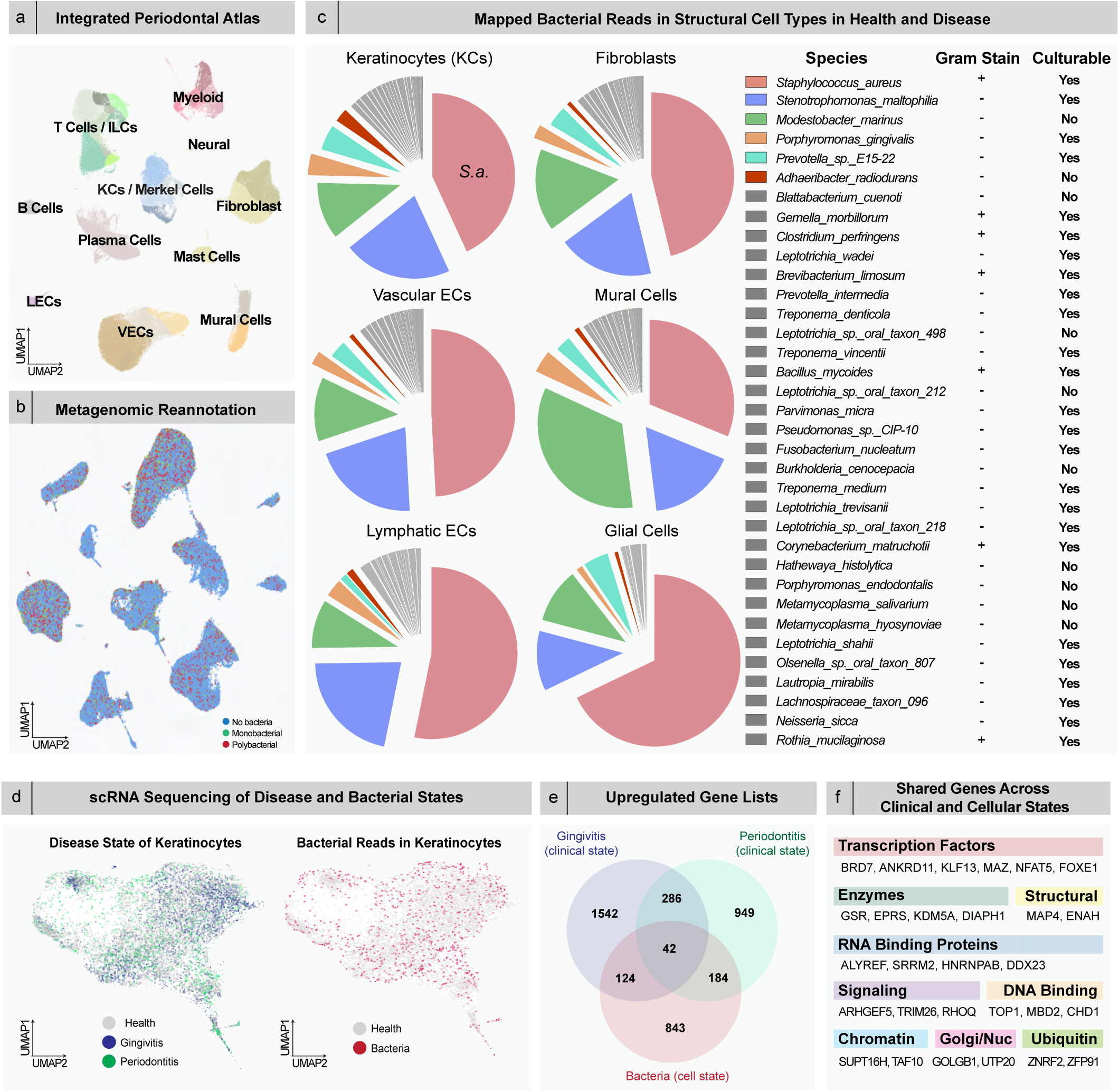
Intracellular bacterial signatures across periodontal cell types. a) We previously generated an integrated periodontal atlas comprising scRNAseq data from 4 independent studies. b) We performed metagenomic reannotation on this atlas using unmapped reads that were mapped back to bacterial sequences. All cell types were classified as having no or containing one (monobacterial) or multiple (polybacterial) intracellular bacterial RNA reads. c) Remapping the bacterial reads across all structural cell types revealed numerous gram-positive and gram-negative species. *Staphylococcus (S.) aureus*, *Stenotrophomonas maltophilia*, *Modestobacter mariunus*, *Porphyromonas (P.) gingivalis*, and *Prevotella* sp. were the most frequently mapped. d) Considering disease states of only keratinocytes (health, gingivitis, and periodontitis), cells contained no, monobacterial and polybacterial intracellular bacterial reads. e-f) Bacteria as a cell state and gingivitis and periodontitis as clinical states all had impacts on gene upregulation, but all 3 independent states shared 42 upregulated genes, including transcription factors, enzymes, RNA and DNA binding proteins, and structural, signaling, chromatin, Golgi/nuclear, and ubiquitin markers. Abbreviations: Keratinocytes (KCs); Innate Lymphoid Cells (ILCs); Lymphatic Endothelial Cells (LECs); Vascular Endothelial Cells (VECs).

This metagenomic reannotation revealed a range of bacterial signatures across all cell types: some cells had no detectable bacterial RNA, others mapped to a single bacterial species (monobacterial), while some displayed more than one species*(20)* (polybacterial) (Figure 1b). All cell types showed at least some bacterial reads. Among structural cells alone, 35 unique species were identified, with 20% being Gram-positive and 80% Gram-negative (Figure 1c). We then assessed the consistency of species proportions across all structural cell types. *Staphylococcus (S.) aureus* (Gram-positive) and *Stenotrophomonas maltophilia* (Gram-negative) were two of the most frequently represented species, although glial cells had a higher proportion of *S. aureus* and *Modestobacter marinus*. Notably, we also found the well-known periopathogen *P. gingivalis* among the most represented species, along with other Gram-negative periopathogens such as *Prevotella intermedia*, *T. denticola*, and *Fusobacterium nucleatum*. Of the list, the majority are known to be culturable*(31)*.

Given the extensive range of mucosal-microbial interactions to analyze within the scRNAseq atlas, we focused on the barrier epithelium, specifically oral keratinocytes across three clinical states (Figure 1d). We identified both mono- and polybacterial signatures in keratinocytes. The bacterial load per cell did not increase with disease (Supplemental Figure 1a,b). However, while the top 10 mono- and polybacterial signatures comprised 56.0% of those detected in health, they accounted for 71.7% in disease, with *P. gingivalis* present only in the disease signatures (Supplemental Figure 1e) and *S. aureus/P. gingivalis* the only unique disease pairing. Given the dysbiotic nature of periodontitis, we examined whether bacterial-associated keratinocytes overlapped with those from gingivitis and/or periodontitis (Figure 1e). We identified distinct signatures for each condition but also discovered a shared transcriptional signature of only 42 upregulated genes out of 3,334 transcripts (∼1%; Figure 1e). Key genes such as *BRD7*, *KLF13*, *MAZ*, *NFAT5*, and *FOXE1* are involved in transcriptional regulation, supporting stress responses in disease*(32)*. Additionally, *ARHGEF5*, *TRIM26*, and *RHOQ* are involved in small GTPase-mediated signaling, relevant to immune responses*(33)* (Figure 1f). Pathway analysis of bacterial-associated keratinocytes revealed significant enrichment in molecular functions like protein binding, oxidoreductase activity, and RNA binding. Biological processes included oxidative phosphorylation, intracellular transport, cell proliferation regulation, and immune response, highlighting how bacteria uniquely modulate keratinocyte function and potentially disrupt epithelial integrity during mucosal inflammatory diseases.

### Polybacterial intracellular macromolecules (PIM) predominate within the host cytosol *in vitro*

Although sparse, the detection of numerous bacterial reads in the mucosal atlas was unexpected, though the increase in periopathogens during disease states was anticipated. The assay did not differentiate whether these changes in keratinocytes were due to cell surface-adhered bacteria or intracellular, host cell-associated bacterial fragments (Figure 1e,f; Supplemental Figure 1d). To investigate the origin of this phenomenon, we cultured primary human oral keratinocytes (HOKs) through several passages. Using a proprietary in-house system for creating spot-based microarrays (<u>M</u>ultiomic <u>A</u>nalyses Thorough <u>P</u>atterned <u>CELL</u> Microarrays, MAPCELL; see Methods), we affixed passaged HOKs to poly-L-lysine-coated slides. In passage 2 HOKs, fluorescence *in situ* hybridization (FISH) revealed that nearly all cells contained intracellular bacterial *16S* rRNA, *fimA* mRNA from *P. gingivalis*, and *fadA* mRNA from the *Fusobacterium* genus (Figure 2a,b). These cells, from two distinct donors, were immediately assayed upon receipt without any manipulation, culture, or additional passaging. Intriguingly, both *KRT19*^hi^ stem/progenitor and *KRT19*^lo^ differentiated HOK subpopulations showed intracellular *16S* signals. This led us to explore whether these HOKs harbored intact intracellular periopathogens.

**Figure 2.**
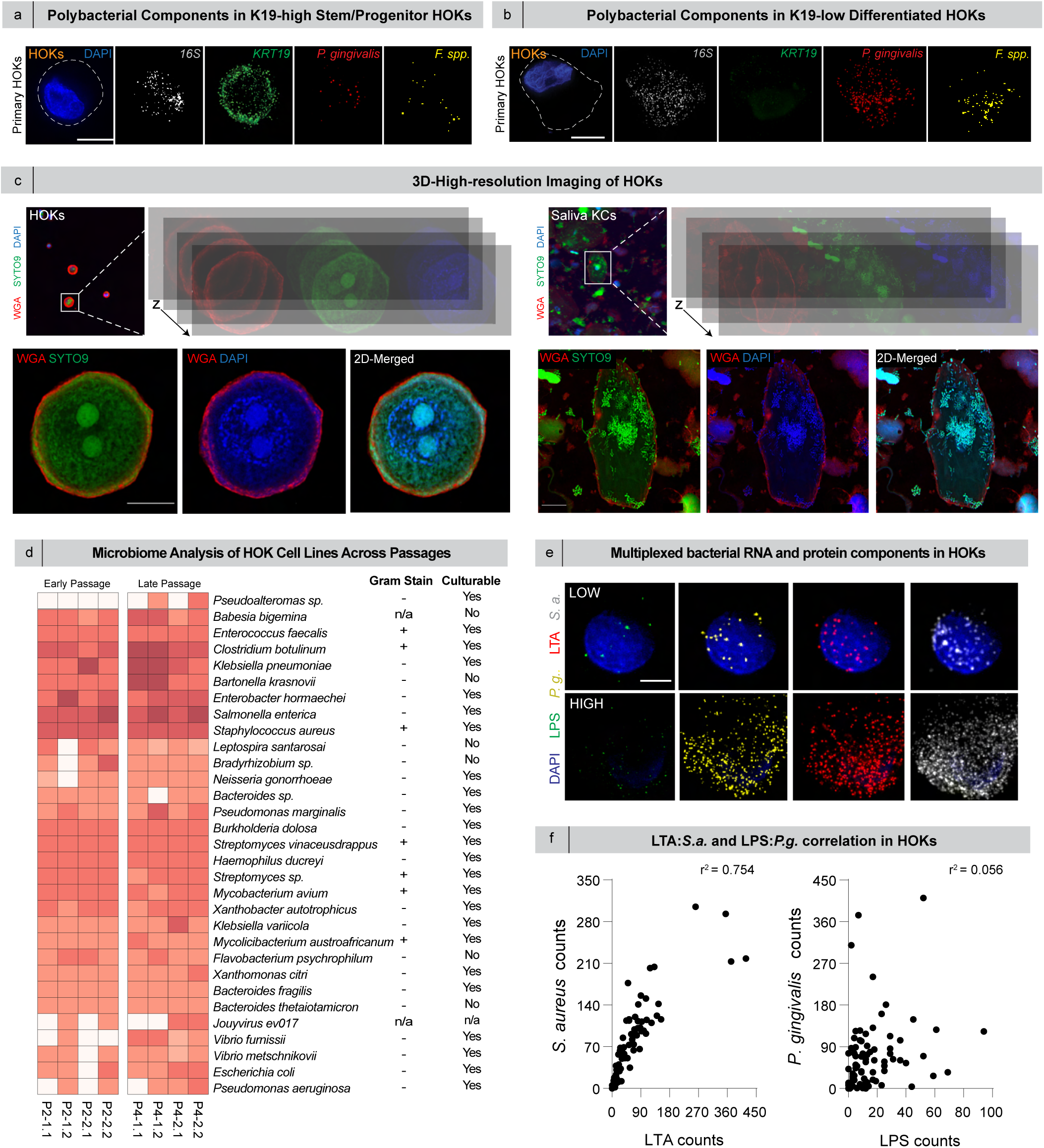
Human gingival keratinocytes (HOKs) contain bacterial rRNA signal, maintained over multiple passages. a-b) RNAscope on P2 HOKs detected bacterial rRNA in a) basal and b) suprabasal human oral keratinocytes. c) We employed 3D super-resolution imaging to detect intact bacteria in P2 HOKs and in saliva. 3D reconstruction of wheat germ agglutinin (WGA) and SYTO9 staining of HOKs revealed no intracellular intact bacteria. Staining of saliva samples, on the other hand, revealed bacteria on the surfaces of cells. d) Microbiome analysis of HGKs, considering as-received (passage [P]2) and multiple passages (P4) thereafter revealed the HOKs contained similar proportions of gram-positive and gram-negative bacterial species across passages. e) Multiplexing anti-lipoteichoic acid (LTA) and anti-lipopolysaccharide (LPS) antibodies with bacterial rRNA probes *fimA* (*P. gingivalis*) and *spa* (*S. aureus*) revealed intracellular bacterial components were maintained at P4. f) LTA and *S. aureus spa* counts had a quasilinear relationship compared to LPS and *P. gingivalis fimA*. Scale bars: a-b, 5 µm; c, 10 µm; e, 5 µm.

We hypothesized that the source of the intracellular rRNA and mRNA signal was intact intracellular bacteria. To investigate this, we used three-dimensional, high-resolution confocal microscopy on passage 2 and 4 HOKs with a dual-staining approach using nucleic acid dyes DAPI and SYTO9, alongside wheat germ agglutinin (WGA) as a membrane marker. DAPI binds to the minor groove of DNA, while SYTO9 intercalates into the major groove, allowing for thorough detection of bacterial DNA fragments or oligonucleotides, regardless of integrity. However, imaging failed to reveal any intact bacteria within the P4 HOKs (Figure 2c; P2 HOKs, Figure 2Sa). As a control, we applied the same approach to shed keratinocytes from human saliva. In contrast to HOKs, saliva-resident keratinocytes showed numerous naturally-occurring bacteria on the cell surface but none intracellularly; these bacteria stained positive for both SYTO9 and DAPI, which made them discernible based on size, shape, and arrangement (Figure 2c). To further validate these findings, we employed transmission electron microscopy (TEM) to search for intracellular bacteria in the HOKs. We observed multiple structures consistent with the breakdown of bacterial cell membranes, not intact bacteria inside autophagosomes, identified by dense 0.5–2.0 µm inclusions within double-membrane, “lasso-like” structures (Figure 2Sb).

Given that both Gram-positive and Gram-negative species had been detected in other assays, we aimed to confirm their presence in HOKs through whole-genome sequencing of two different HOK donor cell lines, at both passages 2 and 4. The microbiome analysis of the keratinocytes revealed bacterial-associated signals across all passages (Figure 2d; Supplementary Data 1). Notably, *S. aureus* was again one of the most represented species, aligning with the scRNAseq atlas data. Additionally, we identified a range of Gram-positive bacteria, such as *Clostridium botulinum*, as well as Gram-negative species, including *Pseudomonas* sp. and *Escherichia coli*. These findings suggest that the signals detected via sequencing and imaging likely originate from bacterial macromolecules, such as rRNA, mRNA, and glycolipids, rather than intact bacteria.

Since HOKs exhibited multiple intracellular bacterial RNA signals over several passages, we aimed to investigate the origin of these signals. We developed custom FISH probes targeting bacterial mRNA (*P. gingivalis*: *fimA*; *S. aureus*: *spa*) and multiplexed them with antibodies (IHC) against lipoteichoic acid (LTA) and lipopolysaccharide (LPS) to differentiate Gram-positive and Gram-negative components within the HOK cytosol (see: Methods). The antibodies were tested on pure cultures of intact bacteria, confirming the anti-LPS antibody reacted with gram-negative bacteria such as *P. gingivalis* (Figure 2Sc), and the anti-LTA antibody with gram-positive bacteria such as *Streptococcus mutans* (Figure 2Sd). As expected, we observed aggregates of LPS^+^ bacteria (Figure 2Se) and LTA^+^ bacteria (Figure 2Sf) displaying known bacterial morphologies on the membranes of salivary keratinocytes. At different passages, HOKs showed varying concentrations of LPS and LTA (Figure 2e), with some cells containing both LPS and LTA inclusions. We also detected mRNA signals for both *P. gingivalis* and *S. aureus* within the same cell using ISH. Plotting the number of LTA counts against *spa* counts revealed a linear correlation (R^2^=0.754); however, no clear correlation was observed between LPS and *fimA* (R^2^=0.056; Figure 2f). This was expected as *P. gingivalis* was not the predominant Gram-negative species in other assays (Figure 1,2). In contrast, *S. aureus* was the most abundant Gram-positive species in both the scRNAseq and whole genome sequencing data, aligning with our FISH findings. This led us to conclude that many observed inclusions were bacterial components. We termed these polybacterial intracellular macromolecules (PIMs), encompassing rRNA, mRNA, glycolipids, and other undetected elements from diverse bacterial origins within the cytosol.

### The concentration of PIMs drives single-cell proinflammatory phenotypes of mucosal epithelia

To further understand the effects of PIMs on mucosal epithelial cells, we used CellChat*(34)* to analyze receptor-ligand interactions within our scRNAseq atlas*(20)*, focusing on how keratinocytes alter signaling patterns based on bacterial presence (see Figure 1). We compared these patterns between healthy and diseased states (Figure 3a). Cell-cell communication patterns were analyzed in keratinocytes with and without bacterial presence and also between healthy and periodontitis states (Figure 3a). Generally, keratinocytes without bacterial presence showed relatively less signaling during periodontitis. However, even in health, keratinocytes with bacteria exhibited broader signaling interactions, particularly with lymphatic and vascular endothelial cells, compared to keratinocytes without bacterial signals (Supplemental Figure 3a,b). In periodontitis, there was a noticeable reduction in signaling between keratinocytes and structural cell types, including fibroblasts and endothelial cells, particularly in keratinocytes lacking bacterial signals. Keratinocytes with bacterial presence in periodontitis maintained some communication with immune cell types, but the signaling intensity decreased, highlighting the potential role of bacterial presence in modulating keratinocyte interactions under both health and disease conditions.

**Figure 3.**
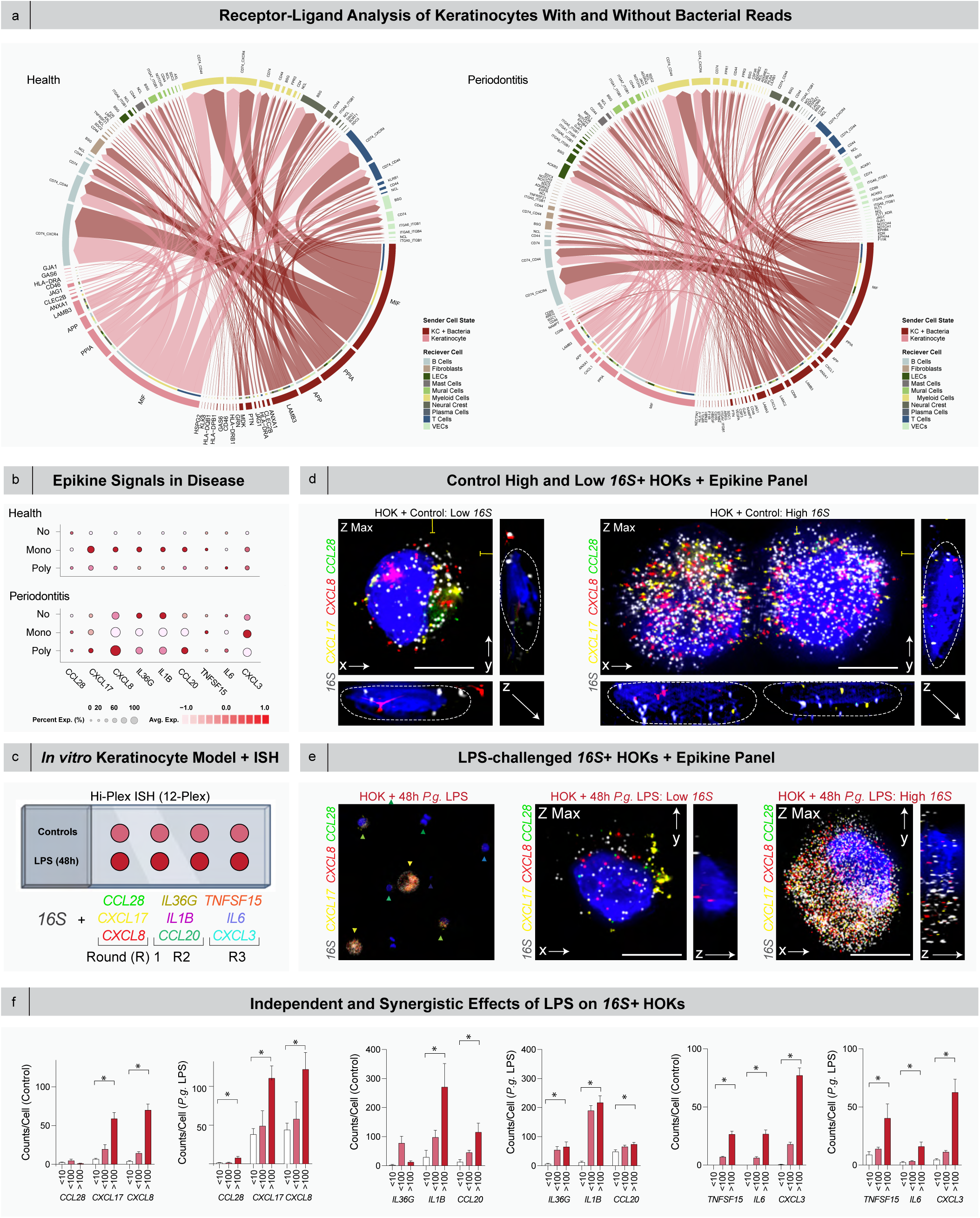
Intracellular bacterial signal affects keratinocyte signaling and correlates with intracellular *16S* signal and LPS *in vitro* challenge of human gingival keratinocytes (HGKs). a) Receptor-ligand analysis showed that in health and periodontitis, signaling by KCs without intracellular bacterial signal decreased, while signaling by KCs with intracellular bacterial signal remained constant. The breadth of signaling to by KCs with intracellular bacterial to LECs and VECs expanded. b) In health and periodontitis, states with no bacteria, monobacterial, and polybacterial signals impacted epikine expression: polybacterial signatures resulted in a shift towards *CXCL17*, *CXCL8*, *IL36G*, *IL1B*, and *CCL20*; only *CXCL3* was a hallmark of monobacterial signature. c) P4 HOKs were cultured with and without LPS (*P. gingivalis*). These cells were assessed for *16S* signal and epikines predicted to increase in disease states using RNA FISH in three rounds. d) HOKs were imaged in xyz using Nyquist-optimized parameters to show correlation of *16S*+ burden and epikine production. This happens without *Porphyromonas gingivalis* LPS (d-e) yet increases in HOKs with LPS, highlighting this phenomenon for *CCL28, CXCL17,* and *CXCL8.* (f) Quantification of all cytokines considering 16S signals per cell. For *CXCL17, CXCL8, IL36G, IL1B, CCL20, TNFSF15, IL6,* and *CXCL8,* intracellular bacterial signature elicits mRNA expression without LPS challenge. Only *CCL28*, which is known to be chemoattractant to T and B cells, is downregulated with increasing bacterial burden. Binning cells in *16S* groups with and without LPS challenge, *CXCL17* and *CXCL8* expression demonstrate a synergy between LPS and polybacterial co-infection. Scale bar: d) and e), 10 μm. p<0.005, paired Student’s T-test (f).

We reanalyzed the epikine data, using bacterial presence as a metavariable. In health, keratinocytes with monobacterial intracellular signals showed upregulation of *CCL20*, *CXCL8*, *CXCL17*, *IL1B*, *IL36G*, and *TNFSF15*, while *IL6* was specifically linked to polybacterial signals. In periodontitis, *CCL20*, *CXCL8*, *CXCL17*, *IL1B*, *IL6*, and *IL36G* were upregulated in keratinocytes either without bacteria or with polybacterial signals, whereas *TNFSF15* remained unchanged, and *CCL28* shifted to mono- and polybacterial states. *CXCL3* was predominantly expressed in monobacterial states but showed higher expression levels across more cells overall (Figure 3b). To validate these findings *in vitro*, we cultured HOKs with varying levels of PIMs. Once P4 control cells reached nearly 50% confluency, we challenged them with 20 µg/mL of LPS for 48 hours. Cells were then fixed, captured and analyzed using MAPCELL. We performed FISH on the same epikines identified in the scRNAseq analysis, utilizing a 9-probe ISH panel (*CCL20, CCL28, CXCL3, CXCL8, CXCL17, IL1B, IL6, IL36G, TNFSF15*; Supplementary Data 2) with *16S* as a reference probe, allowing us to correlate PIMs (*16S*) with epikine expression for each of the three multiplexed rounds (Figure 3c).

We simulated an external challenge to the HOKs by adding LPS to the media. Unchallenged HOKs showed varying *16S* levels, with corresponding differences in epikine expression (Figure 3d; Supplemental 4a). We identified both epikine-low and epikine-high cells with *CCL28*, *CXCL17*, and *CXCL8* expression levels directly correlating with *16S* levels, like patterns seen with LTA (Figure 3d). Upon LPS challenge, there was a clear, albeit non-uniform, shift toward a proinflammatory HOK state. Epikine-low and epikine-high cells were again observed, but the per-cell counts of *CXCL17* and *CXCL8* were significantly higher, again correlating with *16S* levels (Figure 3e). Similar trends were seen for other epikines, including *CCL20*, *CXCL3*, *IL1B*, *IL36G*, *IL6*, and *TNFSF15*.

We then quantified intracellular signals to assess the effect of LPS challenge on *16S+* HOKs. In control HOKs, *16S* alone led to significant upregulation of *CCL20*, *CXCL3*, *CXCL8*, *CXCL17*, *IL1B*, *IL6*, and *TNFSF15*. Following LPS challenge, *CXCL8* and *CXCL17* showed the strongest correlation between LPS exposure and an increased number of cells with epikine expression >100 counts/cell (Figure 3f). Additionally, we found a direct linear correlation between *16S* counts and *CXCL8* and *CXCL17* expression per cell. Apart from *IL1B*, which exhibited a quasilinear trend, no other epikine displayed a linear correlation (Supplemental Figure 4b). These signatures suggest that LPS can synergistically amplify epikine responses, particularly for *CXCL8* and *CXCL17* in cells with few *16S* signals (see: Supplemental Figure 4b). These two chemokines are known to attract neutrophils and other immune cells like monocytes*(35, 36)*. The strong correlation of *CXCL8* and *CXCL17* with *16S* counts in controls, and the shift towards a proinflammatory state in cells with PIMs, indicated that the relationship of keratinocytes in disease with and without PIMs is more complex than currently understood.

### Bacterial association site-agnostically drives proinflammatory tissue phenotypes and pathways

The investigation of HOKs and the *in vitro* LPS challenge model demonstrated that proinflammatory cell states were present even in healthy conditions, but these states could be exacerbated by external factors. Previously, we showed that junctional and sulcular keratinocytes—regionally specialized cells within the crevicular epithelium—were predisposed to proinflammatory phenotypes, though their origins remained unclear. To explore whether these proinflammatory PIMs were a broader phenomenon in both keratinized and non-keratinized mucosal epithelia, we applied the best-correlated epikine panel (*16S*, *CXCL17*, *CCL28*, and *CXCL8*) to periodontitis tissues, examining the site-agnostic effects of *16S* on epikine signaling. Across four mucosal regions—the junction, sulcus, marginal transition zone, and attached gingiva—we identified both *16S*^low^ and *16S*^high^ keratinocytes. In all cases, *16S^l^*^ow^ keratinocytes exhibited minimal epikine expression, while *16S*^hi^ cells upregulated *CXCL17*, *CCL28*, and *CXCL8* (Figure 4a). In each region and type of mucosal epithelium, we binned cells in groups of less than 5 counts and greater than or equal to 5 counts of *16S* rRNA. We then plotted the *16S* ribosomal RNA counts per cell with the epikine counts per cell for these categories (Figure 4b). In the less than 5-count category, we observed little correlation of *16S* with epikines, regardless of region. In the greater than or equal to 5-count category, we observed region-agnostic correlation of *16S* rRNA with each of the 3 epikines; we observed statistically significant higher levels of cells with greater than or equal to 5 counts of *16S* per cell, matching our HOK experiments.

**Figure 4.**
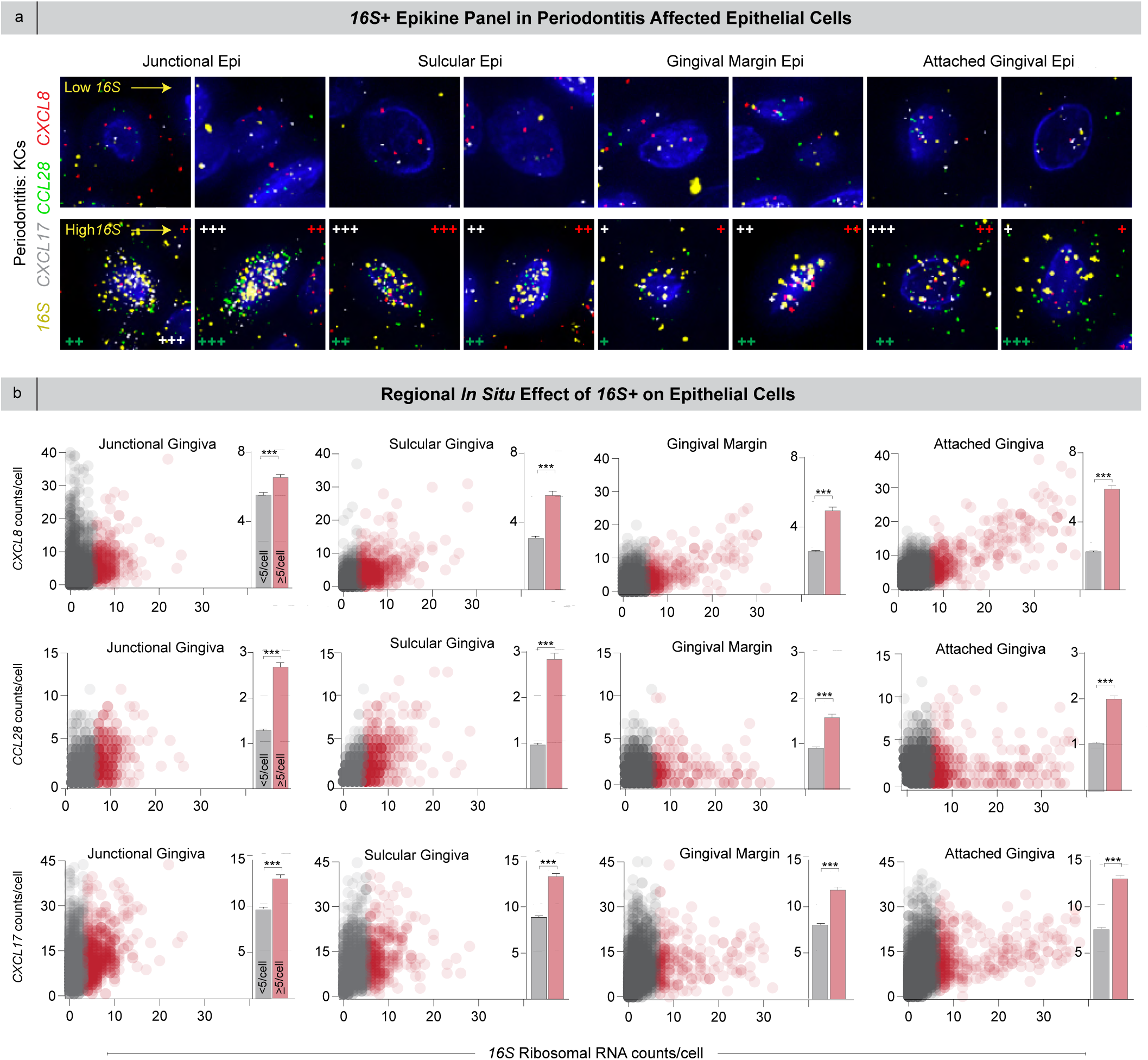
Intracellular bacterial signal *in situ* correlates with epikine signal in periodontitis. a) We used our RNAscope HiPlex epikine panel on human periodontitis biopsies and imaged the junctional, sulcular, gingival margin, and attached gingival epithelial regions. Both low- and high-*16S* keratinocytes were observed in all regions of the epithelium. At a single-cell level, *16S* signal correlated with epikine upregulation, shown here for *CXCL17*, *CCL28*, and *CXCL8*. b) Across the gingival epithelial regions, quantification of intracellular *16S* signal revealed statistically significant associations between bacterial burden and *CXCL17*, *CCL28*, and *CXCL8*. c) Pathway analysis between keratinocytes unassociated with and associated with bacteria reveal significant changes tissue integrity. Scale bars: a) 10 µm. b) p<0.005, paired Student’s T-test.

We then performed pathway enrichment in keratinocytes with and without bacterial association in both health and periodontitis to understand a more global impact of this phenotype *in situ* (Figure 4c). This analysis highlights key pathway shifts in keratinocytes with bacterial association. In health, keratinocytes with bacterial presence show significant enrichment in pathways related to structural and immune maintenance, including COLLAG, CD99, APP, LAMININ, CLEC, PTN, GAS, and CD46. These pathways are crucial for maintaining epithelial barrier integrity, mediating cell adhesion, and supporting immune surveillance. In periodontitis, bacterial-associated KCs exhibit a shift toward pathways that contribute to chronic inflammation and tissue degradation, such as LAMININ, PTN, CD46, EGF, ADGRA, EPHB, MPZ, and CSF3. These pathways suggest enhanced extracellular matrix remodeling (LAMININ), immune response activation (CSF3), and cell migration (EGF, ADGRA), all of which drive the inflammatory cascade and tissue breakdown characteristic of periodontitis. The upregulation of CD46 in both health and disease emphasizes its dual role in protecting mucosal integrity while potentially amplifying immune responses under inflammatory conditions. This suggests that keratinocytes associated with PIMs—here mRNA and rRNA—are uniquely changed within the mucosal epithelial microenvironment.

### The concentration of PIMs drives proinflammatory phenotypes of HOKs and mucosal epithelia

Our analyses suggested that PIMs also drive proinflammatory epithelial phenotypes *in vivo*, though we had only demonstrated this correlation with RNA. To explore the role of bacterial glycolipids and distinguish between Gram-positive and Gram-negative influences, we used MAPCELL to array HOKs onto slides and performed multiplexing with anti-LPS and anti-LTA antibodies alongside *CXCL8* and *CXCL1*—an innate immune epikine highly correlated with *16S(20)*. We identified cells with varying levels of LPS and LTA, often co-occurring within the same cell. Like bacterial rRNA, cells with higher concentrations LPS and LTA levels were frequently *CXCL1*^hi^ and *CXCL8*^hi^ (Figure 5a). We then quantified *CXCL1* and *CXCL8* expression relative to LTA and LPS levels to determine any correlation between epikines and bacterial glycolipids. LTA displayed a direct linear correlation with both *CXCL1* and *CXCL8* (*CXCL1*: R^2^=0.95; *CXCL8*: R^2^=0.97), while LPS showed a broader range with weaker correlation to these epikines (*CXCL1*: R^2^=0.32; *CXCL8*: R^2^=0.34; Figure 5b). Unchallenged HOKs from two separate lots exhibited the same LTA *CXCL1/8* correlation and lower, but better, correlation with LPS (Supplemental Figure 5a). In additional LPS-challenged primary cells, similar LTA/LPS trends were observed, with LTA more strongly correlating with innate epikine expression (Supplemental Figure 5b).

**Figure 5.**
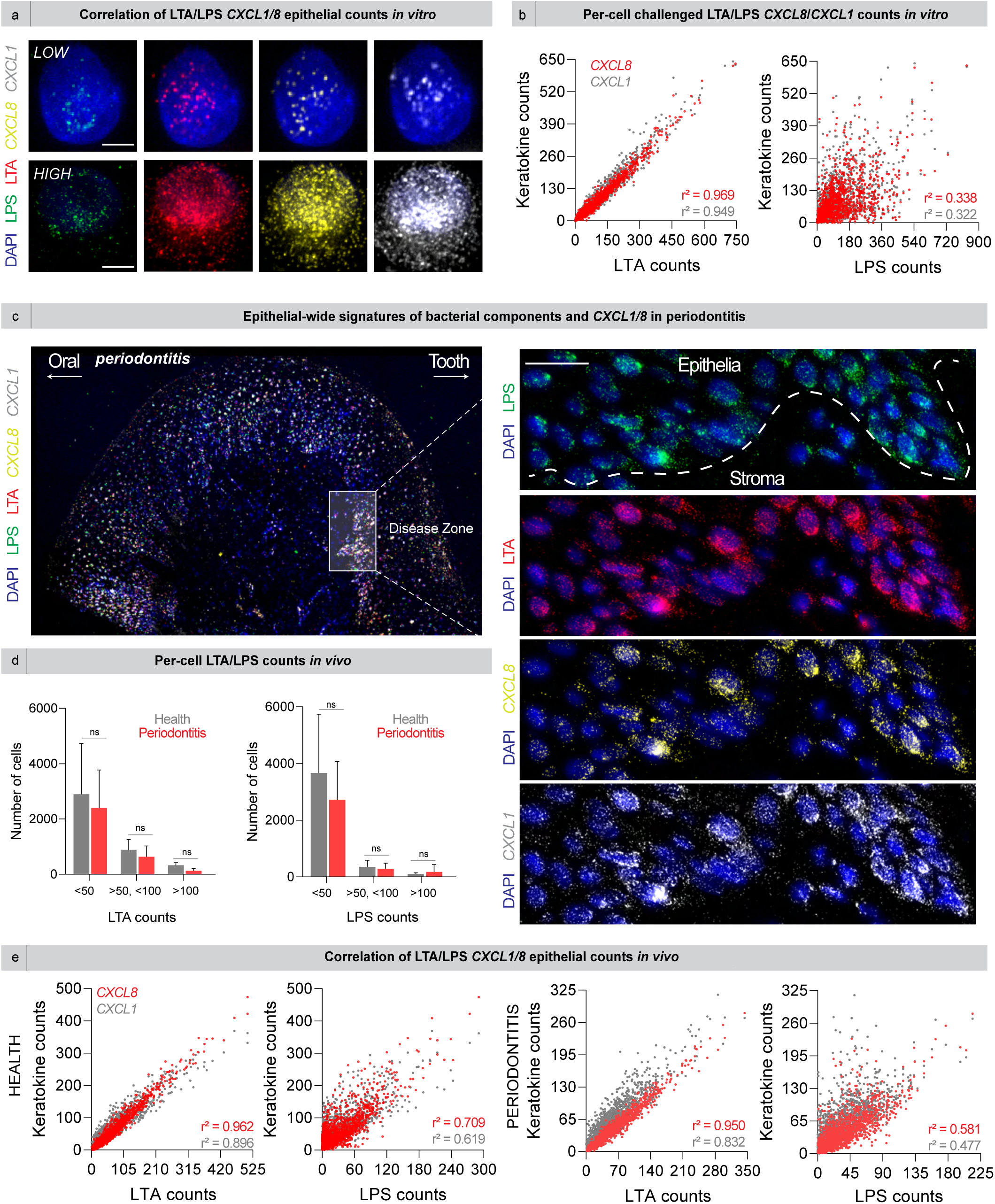
Gram-positive bacterial glycolipids correlate with epikine signaling *in vitro* and *in vivo*. a) Multiplexing anti-LTA and anti-LPS antibodies with epikines *CXCL8* and *CXCL1* in HOKs revealed synergistic upregulation of epikines with increasing intracellular LTA. b) We quantified per-cell correlation of bacterial glycolipids and epikines and found that *CXCL1/8* both increased linearly with increasing LTA counts. On the other hand, epikine counts had little correlation with increasing LPS counts. c) In periodontitis tissues, we observed epithelial-wide signatures of bacterial components and *CXCL1/8*. In the disease zone of the sulcus, we found LPS- and LTA-low and -high cells whose *CXCL1/8* directly correlated with bacterial component presence. d) We binned keratinocytes in health and periodontitis based on their LTA and LPS counts, site agnostic of the epithelium. For both LTA and LPS, regardless of health or disease status, we found no statistically significant difference between the number of cells with less than 50 counts, cells with 50 or greater but less than 100 counts, and cells with greater than 100 counts. e) We quantified the correlation of epikine counts in health and periodontitis tissues with intracellular LTA and LPS counts. In both health and periodontitis, increasing *CXCL1/8* counts had higher correlation with LTA than with LPS. Abbreviations: lipoteichoic acid (LTA); lipopolysaccharide (LPS). Scale bars: a) 5 µm; c) 20 µm.

We then extended the assay to tissues, this time analyzing both healthy and diseased mucosa (periodontitis, Figure 5c; health, Supplemental Figure 5c). In both states, we detected the presence of LPS and LTA distributed throughout different mucosal epithelial regions and cell types. In the basal stem cell layer of the sulcular epithelium, we observed the co-occurrence of both LPS and LTA within the same basal cells. Like our findings in HOKs, cells could be clearly classified as LPS/LTA-low and -high, though no region showed specificity for these cell types. Notably, the highest epikine expression for *CXCL1* and *CXCL8* was observed in PIM-high cells. With no discernable trends based on region or mucosal type, we applied cell segmentation using a pretrained deep learning algorithm to further investigate (Cellpose 2.0*(37)*; see Methods). In both health and periodontitis, we quantified the number of LTA and LPS counts on a per-cell basis. Cells were binned into less than 50, from 50 to 100, and greater than 100 counts of both LTA and LPS. When we compared health and disease, we found no statistically significant differences between these bins (Figure 5d), mirroring data from scRNAseq (Figure S1a,b). In both health and periodontitis across all samples, we found that intracellular LTA correlated with *CXCL1/8* upregulation; LPS did not have the same correlation, matching our *in vitro* assays (Figure 5e; additional examples, Figure S5e).

### PIMs predictably impact mucosal epithelial cell functions and inflammatory milieu in health and disease

With the intracellular bacterial signatures in hand, we wondered how these signatures were associated with shifts in epithelial cell-cell communication to other cell types, signatures of host-impact outside of epikines, and association with pathways. To better understand this, we implemented CellChat on the scRNAseq data, considering bacterial association with KCs as a metavariable. In both health and periodontitis, myeloid cells had the highest incoming interaction strength, and fibroblasts had the highest outgoing interaction strength. We observed higher incoming interaction strength in KCs with bacterial state than in those without (Figure S6a). Next, we asked whether same- and intercellular signaling strength was impacted. While KCs without bacterial state decreased in signaling strength, KCs with remained relatively unchanged, though we also observed stronger plasma and T cell signaling strength in periodontitis but lower VEC and fibroblast signaling strength (Figure S6b).

We next analyzed bacterial association signatures (none, mono-, or polybacterial) in keratinocytes from both healthy and periodontitis samples (Figure S6c), revealing distinct gene signatures based on bacterial association. In periodontitis, keratinocytes without bacteria shifted from epithelial identity markers like *CXCL14*, *FDCSP*, and *SFRP1*. Keratinocytes with bacterial intracellular signals showed the most significant shift, with over 20 genes exhibiting a log fold 1 increase in expression, including upregulation of *CSF3*, *CXCL1*, *IL1RN*, and *MMP7*—genes previously associated with periodontitis*(38)*. These gene signature changes suggest that bacterial presence can drive toward more inflammatory phenotypes, contributing to disease pathology.

### PIM-induced cellular responses influence clinical and translational outcomes on keratinocytes in health and periodontitis

To assess the clinical and translational relevance of our findings, we applied the Drug2Cell pipeline*(28)* using our integrated scRNAseq atlas (see Methods). Drug2Cell is a computational tool designed to predict drug efficacy based on cellular profiling, though it had not previously been used with meta-annotations like bacterial presence. In this version, Drug2Cell analyzes single-cell RNA sequencing data to examine gene expression profiles and signaling pathways, integrating this with known drug targets to predict how effective specific drugs will be on certain cell populations. The algorithm generated top-predicted drug targets for structural and immune cell types, factoring in bacterial states (Figure 6a; Supplementary Data 2). Targets were classified in two ways: 1) by drug category (keratinocyte-specific, antibiotic, monoclonal antibody, chemotherapeutic, and anti-inflammatory), and 2) whether drugs were shared between bacterial and non-bacterial states or unique to one. Overall, nearly all drugs had higher predicted efficacy in keratinocytes with bacterial associations, both in healthy and periodontitis conditions.

**Figure 6.**
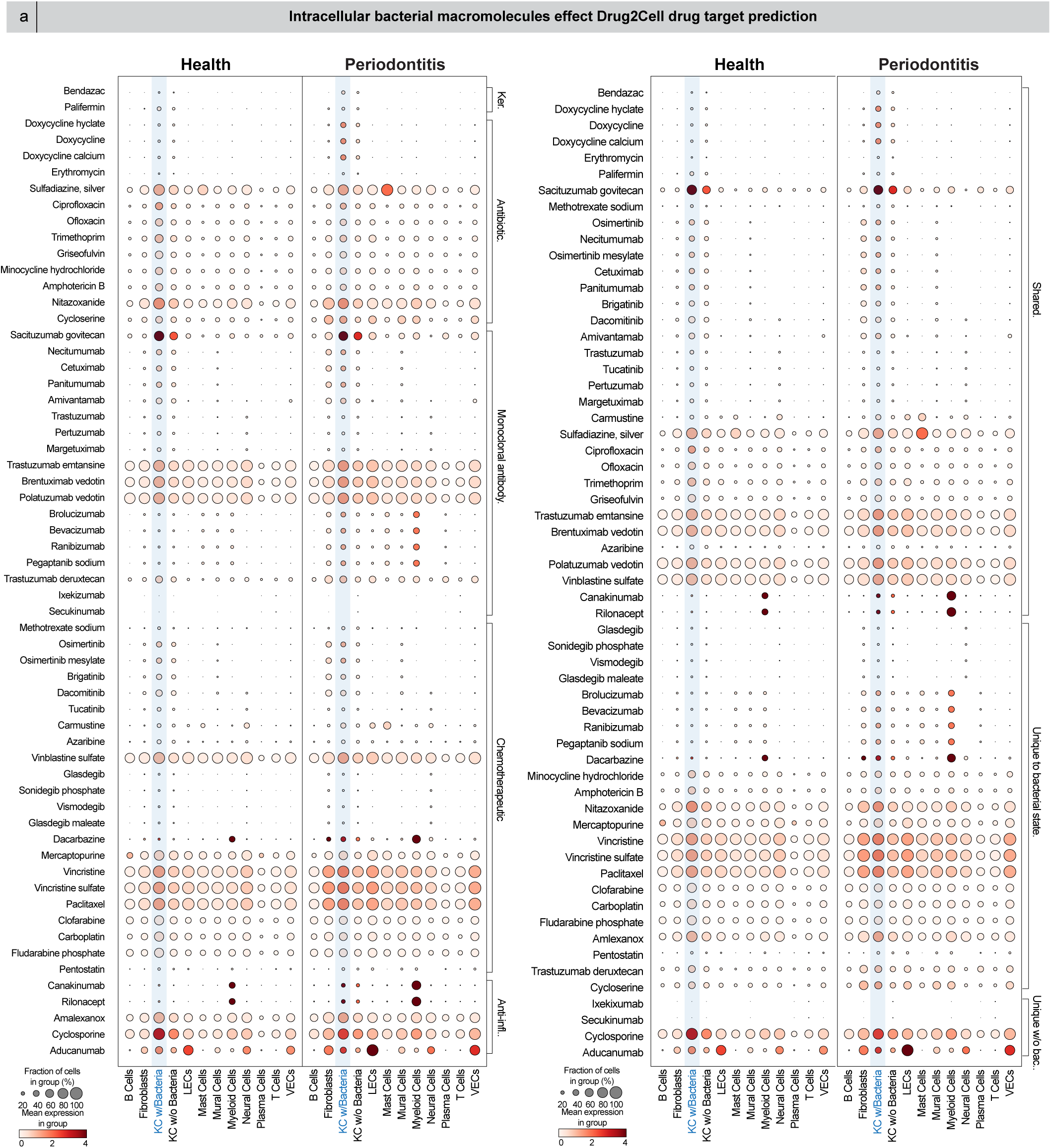
Drug2Cell analysis of the impact of intracellular bacterial components (IBCs) on predicted drug targets. a) Impact of intracellular bacterial components on drug target prediction through Drug2Cell. Drugs were organized by either drug category or whether the drug was shared between or unique to keratinocytes (KCs) with or without bacteria. Drug categories included keratinocyte-specific, antibiotic, monoclonal antibody, chemotherapeutic, or anti-inflammatory. KCs with bacterial signal had higher predicted targets than KCs without. Abbreviations: Keratinocyte-specific (Ker.), Anti-inflammatory (Anti-infl.); rest: see Figure 1.

Some of the shared drug targets included widely used periodontitis treatment doxycycline hyclate, along with other antibiotics such as erythromycin, ofloxacin, ciprofloxacin, and trimethoprim. Drug2Cell also predicted efficacy for monoclonal antibodies and chemotherapy drugs, with key shared pathways including EGFR (e.g., dacomitinib and amivantamab) and HER2 (e.g., trastuzumab and tucatinib) targeting, as well as DNA intercalation pathways (e.g., methotrexate sodium and carmustine). Immunosuppressive treatments like the IL1B-inhibitor canakinumab and keratinocyte-specific drugs such as bendazac and palifermin—commonly used for preventing severe oral mucositis—also emerged as important targets.

While many drugs were shared between keratinocytes with and without bacterial presence, unique drug predictions were also identified. For keratinocytes harboring bacterial signatures, 140 unique drug targets were identified compared to just 4 for keratinocytes without bacterial signals with 95 shared. Predicted treatments for bacteria-positive cells spanned antibiotics (e.g., minocycline, nitazoxanide), anti-inflammatory agents (e.g., amlexanox), chemotherapy drugs (e.g., dacarbazine), and monoclonal antibodies (e.g., trastuzumab deruxtecan). Novel antibody targets included VEGF (brolucizumab and bevacizumab) and sonic hedgehog pathway inhibitors (glasdegib and sonidegib). While some predicted targets emerged for bacteria-negative cells, potential reactivity with bacteria-positive keratinocytes was still noted, emphasizing the complexity of these interactions.

## DISCUSSION

This study highlights the discovery of polybacterial intracellular macromolecules (PIMs) in human upper airway mucosal epithelial cells. We observed this phenomenon in both keratinized and non-keratinized oral mucosal epithelial types, revealing a broad and pivotal role for PIMs in modulating immune responses, particularly in niche-specific oral inflammation seen in conditions such as gingivitis and periodontitis. Through reanalysis of our single-cell RNA sequencing atlas*(20)* and high-resolution microscopy, we uncovered a strong association between Gram-positive bacterial macromolecules and immune activation (epikine) in human oral keratinocyte niches. This finding suggests that these intracellular bacterial components play a substantial role in sustaining inflammation, even after the whole intact bacterium has been cleared by mechanisms like phagocytosis and autophagy. These findings open new paths for understanding host-microbe interactions within mucosal barriers and their consequences for mucosal inflammatory diseases.

Notably, our findings revealed a linear relationship between Gram-positive PIMs and the upregulation of key inflammatory mediators, *CXCL1* and *CXCL8*, which may underlie the persistent inflammatory epikine milieu observed in epithelial cells. This modulation of the immune landscape by Gram-positive bacteria is not limited to oral health. Bacteria-associated single epithelial cells within tumors exhibited increased expression of pro-metastatic molecules, such as *PLAU/R* and *AREG*, along with the same upregulated epikines, *CXCL1* and *CXCL8(39)*. These findings—focused on intact *Fusobacterium nucleatum*—demonstrated how bacteria within microniches could significantly alter host cell signaling, driving immune recruitment and disease progression, including cancer metastasis. This concept of “epikine” signatures expands our understanding of the consequences of microbial presence in epithelial niches, linking chronic inflammation with disease progression beyond the oral cavity. Further, we found that Gram-positive PIMs were broadly expressed across the oral cavity, likely due to the ability of these bacteria to effectively colonize and persist on and within mucosal tissues*(12)*.

A critical aspect of our findings concerns the origin and nature of these PIMs. It is also important to recognize the significant historical contributions that have shaped our understanding of intracellular bacteria in oral tissues. Notably, regarding the microbial composition of crevicular epithelial cells, species such as *P. gingivalis* and *P. intermedius* were more prevalent in the epithelial cell layer compared to unattached bacteria*(40)*. This finding was critical in demonstrating that these pathogenic bacteria were capable of associating closely with epithelial cells, an early observation that suggested bacterial invasion of host tissues might be a key feature in periodontal diseases. Further, various oral bacterial species were identified on or within gingival crevice epithelial cells in chronic periodontitis patients, including species such as *Porphyromonas gingivalis*, *Treponema denticola*, and *Prevotella intermedia(41)*—three species all detected in our single-cell atlas (Figure 1). These findings indicate the persistent invasion and potential for long-term pathogeneic effects within periodontal tissues.

While initially identifying the PIMs as intact bacteria, then recognizing some are not associated with intact bacteria, their precise form remains uncertain. These PIMs could be outer membrane vesicles (OMVs) or other bacterial extracellular vesicles (BEVs), produced by both Gram-positive and Gram-negative bacteria*(42, 43)*. OMVs contain bacterial proteins, endotoxins like LPS and LTA, immunogenic molecules, and other virulence factors*(44)*. When released into the environment, these vesicles can be taken up by host cells through fusion with the host plasma membrane*(45)*, free diffusion*(46)*, or endocytosis*(47)*, where they bypass extracellular immune surveillance and directly activate intracellular immune pathways by delivering immunogenic cargo such as glycolipids (LPS or LTA), RNA, and proteins*(48)*. OMV and BEV bacterial component delivery to host cells could explain the origin of the persistent inflammatory signals we observed in epithelial cells. The cargo within these vesicles can resist intracellular degradation*(49)*, which contributes to intracellular vesicle persistence and cargo engagement with inflammatory response signaling pathways*(50)* and drives immune activation, seen across multiple pathogens *(51)*.

Alternatively, it is possible that some of these PIMs are remnants of bacterial degradation processes within host cells, particularly those mediated by autophagy wherein pathogens and bacterial components may have been digested into smaller macromolecules. The PIMs we identified may be byproducts of this autophagic processing, where the host cell successfully degraded bacterial invaders but retained immunostimulatory LPS, LTA, or bacterial RNA. These remnants may persist within autophagosomes or lysosomes, driving immune activation even after the pathogen has been neutralized. Indeed, bacterial persistence within autophagic vesicles has been reported in other infectious diseases, such as *Mycobacterium tuberculosis*, where bacteria or their components evade complete degradation and perpetuate chronic inflammation*(52)*.

At this stage, we cannot definitively determine whether these PIMs are intact membrane-bound vesicles, like OMVs, or free bacterial macromolecules processed by host autophagic mechanisms *in situ*. To clarify this, future studies using advanced imaging techniques, such as transmission electron microscopy on tissue sections, are necessary, which would allow us to potentially differentiate between intact bacterial vesicles and degraded remnants within autophagosomes or lysosomes, providing critical insights into how bacterial components persist and trigger immune responses in epithelial cells. Understanding the physical state of these PIMs is crucial, as it would clarify whether they remain membrane-bound or exist as free bacterial macromolecules within host cells. Microbes like *S. aureus* via extracellular or intracellular glycolipids may sensitize epithelial cells to invasion and long-term colonization*(53)*. This distinction could reveal the specific mechanisms by which commensal and pathogenic bacteria persist within host cells and contribute to chronic inflammation.

Additionally, while we successfully detected bacterial macromolecules in some cells, there are inherent limitations to our approach. Single-cell RNA sequencing is not designed to comprehensively capture microbial reads; likely, some cells harboring PIMs were missed in our analysis*(30)*. Bacterial component incidental co-capture in single-cell sequencing is challenging, especially in a complex tissue environment like the gingiva. Expanding the resolution of bacterial detection would allow us to capture a more comprehensive picture of how bacterial communities interact with epithelial cells and contribute to chronic inflammation. Further, in this study, we focused primarily on keratinocytes, but other epithelial cell types, such as stromal cells, fibroblasts, or immune cells, may also be affected by bacterial macromolecules. These cell types could have distinct roles in modulating the immune response and the persistence of inflammation. Future studies should explore whether other epithelial or immune cell types also harbor PIMs and how their responses differ from those of keratinocytes. There may also be species-specific effects on cell-cell communication, and different bacterial components may uniquely affect host cells, particularly in the context of live bacteria versus bacterial remnants. It is likely that species-specific variations in bacterial components influence immune responses and the progression of chronic inflammatory diseases.

Despite these limitations, our study provides significant insights into the role of bacterial macromolecules in driving inflammation and immune responses within the oral mucosa. These findings likely reflect a broader phenomenon across barrier epithelial cells, which form the first line of defense against microbial invasion. The persistence of bacteria and their components, whether as OMVs, EVs, or autophagic remnants, may represent a conserved immune strategy that harkens back to ancient host-microbe interactions. Epithelial cells across the aerodigestive tract, including the respiratory and gastrointestinal systems, may similarly retain bacterial macromolecules, contributing to sustained immune activation in chronic inflammatory diseases beyond the oral cavity. Our work underscores the importance of understanding bacterial persistence and the role of PIMs in modulating immune responses. Whether these PIMs are intact bacterial vesicles or the remnants of bacterial degradation, they represent a novel mechanism by which host-microbe interactions can sustain chronic inflammation. Future studies will need to explore how these bacterial components are processed and retained within host cells and how they contribute to disease states across mucosal barriers. Expanding our knowledge of these processes will provide new opportunities for targeted therapies aimed at resolving chronic inflammation and restoring tissue health.

## MATERIALS AND METHODS

### Study Design

The objective of this research was to first understand how bacteria in mucosal epithelial cells drove *in vitro* and *in situ* inflammatory signaling, then to understand how persistent polybacterial intracellular macromolecules (PIMs) in mucosal epithelial cells drove single-cell inflammatory signaling and impacted predicted drug efficiency. Units of investigation were gingival biopsies from healthy and periodontitis volunteers and primary human oral keratinocytes (HOKs) purchased commercially. Observations were made through computational analyses, RNA fluorescence *in situ* hybridization (RNA FISH), and immunofluorescence (IF); data for the latter two were acquired using a Leica DMi8 with THUNDER Imager. Samples for RNA FISH and IF were from n = 3 gingival biopsies for health and periodontitis and n = 2 for HOKs. All generated data were included in analyses; no outliers emerged during this study. Blinding was not possible during collection or analysis because participant allocation into health and periodontitis necessitated knowledge of clinical metadata. Histopathological and fluorescence microscopy analysis of biopsies revealed health or disease status of samples despite blinding where possible to identity. We were blinded during computational analyses.

### Statistical Analysis

All non-sequencing-based data were analyzed in Fiji*(54)*, QuPath (using CellPose 2.0 for segmentation*(37)*), and/or Prism 9. The selection of statistical tests is described in the text and figure legends; all statistical tests were two-sided. The graphs supporting each Figure and Supplementary Figure were generated using Prism 9/10 (GraphPad) and Trailmaker (Parse Biosciences) unless otherwise specified.

## Acknowledgements

We acknowledge Victoria J. Madden, Jillann A. Madren, and Kristen K. White in the Microscopy Services Laboratory, Department of Pathology and Laboratory Medicine, UNC Chapel Hill for TEM imaging.

## Author contributions

Conceptualization: KMB, QTE, ZAM

Methodology: QTE, ZAM, BTR, TW, AT, SHR, KIK, KMB

Resources: KMB, TW, AT, NSO, AH, ANG, AJK, PRT, SHR, HK, KIK

Investigation: QTE, ZAM, MK, BFM, BTR, ZR, JC-P, SHR

Visualization: QTE, ZAM, MK, ZR, JC-P

Funding Acquisition: ZR, KMB

Project administration: QTE, ZAM, KMB

Supervision: QTE, ZAM, KMB

Writing – original draft: KMB, QTE, ZAM

Writing – review & editing: KMB, QTE, ZAM, MK, BFM, BTR, TW, ZR, AT, JC-P, AH, ANG, AJK, PRT, SHR, HK, KIK

## List of Supplementary Materials

Materials & Methods

Figure S1 to Figure S6

Supplementary Data S1 and S2

Code/Scripts (http://github.com/loci-lab/periodontitis)

## MATERIALS AND METHODS

## ETHICS STATEMENT

This research complies with all relevant ethical regulations. Studies using human gingival biopsies were approved by the University of Pennsylvania (IRB #6; Protocol #844933; Lead PI: KIK). Studies using human saliva and human nasal and tonsil samples were approved by the University of North Carolina at Chapel Hill (human saliva, IRB #24-0944, human nasal samples, IRB #22-1786; Lead PI: AJK), and Duke University (IRB #Prooo114526; Lead PI, PRT), respectively.

## HUMAN INTEGRATED PERIODONTITIS ATLAS

### Database generation and subclustering

We have previously generated a “V1” integrated periodontitis scRNAseq atlas, the full details of the generation of which can be found elsewhere. The publicly available version of this atlas can be found at https://cellxgene.cziscience.com/collections/71f4bccf-53d4-4c12-9e80-e73bfb89e398. The scRNAseq dataset was processed, analyzed, and visualized using Trailmaker, currently hosted by Parse Biosciences (formerly known as the Cellenics® community instance hosted by Biomage and initially accessed between May 2022 and February 2024).

## HUMAN PRIMARY CELL CULTURE AND ANALYSIS

### Primary cell culture of (HOKs)

All reagents were purchased and used as received from Lifeline Cell Technology unless otherwise noted. Cells were plated, passaged, and stored based on our lab’s cell culture protocol. In brief, passage (P)2 HOKs (product #FC-0094, lot #05390 and #10781) were thawed, aliquoted into 100 µL portions, and diluted to 1.5 mL in fully supplemented DermaLife K Keratinocyte Medium. The P2 HOKs were added to 6-well Nunc plates (Thermo Fisher) in these aliquots and grown at 5% CO_2_ and at 37°C. The media was changed after 24h, then 48h thereafter until cells reached a minimum of 70% confluency. The P2 HOKs were then passaged using 0.5% trypsin-EDTA, neutralized, and pelleted using a Sorvall Legend X1R centrifuge (Thermo Scientific) at 1232 RCF for 5 min, resulting in P3 HOKs. Some P3s were then replated and grown the same procedure to obtain P4 and P5 HOKs.

### HOK cryopreservation and fixation

In-house passaged HOKs that were not used in experiments were either cryopreserved or fixed. For cryopreservation, the HOKs were centrifuged as above, the supernatant was removed, and the HOKs were resuspended in 1 mL of Frostalife. HOKs were cooled to -80°C in an ultra-low-temperature freezer for at least 2h using a Nalgene Freezing Container before immersion in liquid nitrogen. For fixation, following centrifugation and removal of the excess solution, the cells were suspended in 4% paraformaldehyde (PFA) for a minimum of 24h. Following fixation, the PFA solution was removed, and the cells were resuspended in 70% ethanol (EtOH) in water. These were stored in a 4°C refrigerator until future use.

### Lipopolysaccharide (LPS) challenge of HOKs

LPS from *Porphyromonas gingivalis* was added to endotoxin-free water (Invivogen; #tlrl-pglps). Serial dilution was used to prepare a solution of 20 µg/mL of *P. gingivalis* LPS in HGK medium (LPS medium). P3 HOKs were plated in a 6-well plate and grown until a minimum of 40% confluency. Then, the HOK medium was removed, and 1.5 mL of LPS medium was added to three of the wells; the other three wells were filled with standard HOK medium. The cells were grown for 48h before the medium was removed. The cells were passaged and either cryopreserved or fixed.

### Mycotesting of HOKs

Mycotesting was performed using a previous protocol from our lab*(20)*. All cells were found to be mycoplasma negative.

### DNA isolation for sequencing

The DNA isolation protocol for sequencing from HOKs has been previously described*(55, 56)*.

### Illumina whole genome shotgun (WGS) sequencing

The full experimental details for WGS sequencing performed on HOKs and culture media can be found elsewhere*(57, 58)*.

### Transmission Electron Microscopy (TEM) of HOKs

The full experimental details for TEM of HOKs are described elsewhere*(59)*. Samples were viewed using a JEM-1230 (JEOL) transmission electron microscope operating at 80 kV (JEOL USA, Inc., Peabody, MA) and images were obtained using a Gatan Orius SC1000 CCD Digital Camera and Gatan Microscopy Suite 3.0 software (Gatan, Inc.).

## VALIDATION OF RNA AND PROTEIN IN CELLS AND TISSUES

### Tissue preparation, mounting, and sectioning

Deidentified human gingival tissues from gingival biopsies (UPenn to LOCI; IRB#844933; MTA#68494) were placed in 10% N-buffered formalin (NBF; Sigma) and fixed for a minimum of 24h at 4°C, washed in PBS, then placed in 70% ethanol (EtOH) until embedding in paraffin blocks using a Leica system. These formalin-fixed, paraffin embedded (FFPE) tissues were sectioned using RNAse precautions on a Leica system onto SuperFrost Plus slides (Fisher Scientific).

### Preparation of Poly-L-lysine coated slides

The following steps were conducted in a biosafety cabinet to reduce the adhesion of dust particles to the slide. SuperFrost Plus slides (Fisher, 12-550-15) were dipped in 70% ethanol, wiped dry with a dust-free paper towel, and fully air dried. In a Coplin jar, a stock solution of 0.01% Poly-L-lysine solution was prepared by diluting 0.1% Poly-L-lysine solution (Sigma, P8920-100) 1:10 with Invitrogen RNAse-free sterile distilled water (Fisher, 10-977-015). The Coplin jar of 0.01% Poly-L-lysine solution was capped and pre-warmed in a 37°C incubator for 1h. Up to 5 clean slides were loaded into a Five Slide Holder (Electron Microscopy Sciences, 7140-06); then, this assembly was snapped into a clean Coplin jar lid. The stock solution was removed from the incubator, and the lid was removed inside the biosafety cabinet. The lid containing the slide assembly was gently screwed onto the top of the 0.01% Poly-L-lysine solution and incubated for 30 min at 37°C, after which it was transferred to a clean, dry Coplin jar and dried for 30 min, also at 37°C. In the biosafety cabinet, slides were removed from the Coplin jar, and the slide assembly was rested diagonally on a paper towel to drain and dry in the airflow.

### Preparation of well frames for Multiomic Analyses Thorough Patterned CELL Microarrays, (MAPCELL)

In the biosafety cabinet, well frames were added to the slide on the same day of slide coating. A ceramic digital hot plate (Fisher, S29042) was pre-heated to 200°C. Eight 5-mm clear thermoplastic beads (Perler, 80-19019) were arranged in 2 rows of 4 beads in the center of a coated slide. Beads were affixed to the slide surface using heat/cool cycles at timed intervals. The slide with the beads was carefully placed on the hotplate for 20 s, then removed and cooled for a total of 20 s. For the first 10 s of cooling time, the slide was cooled on a 5 mm stack of paper towels. During the second 10 s of cooling time, the beads were then gently pressed with gloved fingers using equal pressure distribution to seal them to the slide. The process was then repeated two to three times until each bead formed an even circular shape. Blank templates were prepared if needed for centrifuge balance.

### Addition of fixed cells to slides using MAPCELL

To test for cell distribution prior to using the custom cell affixing device, the fixed cell solution was vortexed, and one 5-µL droplet of cells per sample was pipetted onto an uncoated microscope slide. Droplets were deposited in the same pattern as beads in the well-frames. The slide was dried fully at 60°C on a plate warmer or hot plate. To eliminate salt crystal residue from the PBS, the slide was dipped thrice in RNAse-free distilled water, then fully dried again at 60°C. Cells were observed under an AxioVert.A1 (Zeiss) microscope at low power to examine distribution on the slide, as this created an approximation of cell affixation to a coated slide using the device. At this time, any adjustments in concentration were made prior to affixing the cells with the device onto the poly-L-lysine coated slides.

In a biosafety cabinet, cells were loaded into the previously prepared well-frames. 5 µL of suspended cell solution was loaded per well at a concentration of ∼40 cells per µL. Care was taken to not scratch the bottom of the wells or pipet air into the wells. After the wells were loaded, the well frames were inspected for fluid distribution, checking to ensure no droplets had formed on the sides or bottoms of any of the wells. If any droplets formed, a fresh pipet tip was used to draw up and re-deposit the fluid into the well evenly. Proper distribution was evidenced by an even, thin film of fluid uniformly distributed in each well and was essential for the successful deposition of cells consistently onto the slide.

A stack was created by placing a plastic aerated lid (3D printed with polypropylene, 75 × 25 × 3 mm with 2 × 4 mm tab and 0.2 mm evenly distributed perforations) on top of each loaded well frame on the slide. To assemble a device, each stack was inserted into an adjustable metal clip for glass (Antrader, a170826WQ001) such that the well frame was balanced between the two screws, with the end of the slide extending beyond the balanced area. Each device was secured by tightening two screws on the metal clip by gently alternating between each screw until the assembly was snug. Blank devices containing matching well-frame patterns were assembled at this time to balance the centrifuge. No fluid was added to the blanks.

MAPCELL arrays containing cell solution and blanks were assembled and loaded into custom-built slide centrifuge adaptors (125 mm W × 30 mm T × 85 mm D). These adapters were loaded into a microplate rotor (Eppendorf, A-2-MTP) for a swinging bucket centrifuge (Eppendorf, 5430). The devices were centrifuged at 1,600 RPM for 20 min. After centrifugation, clips were removed from the devices in a biosafety cabinet in the same alternating pattern. Fluid in the well frames was allowed to air dry by placing each slide on a paper towel stack cut to the dimensions of a slide (75 mm L × 25 mm W × 5 mm H) in the back of the biosafety cabinet, next to the air vents. Samples were allowed to dry under the cabinet airflow for approximately 1h; then, the well-frames were removed with tweezers. Slides were baked on a slide warmer at 60°C for 10 min. To remove the salt crystals deposited by the PBS, slides were dipped thrice in RNAse-free distilled water, then dried again on the slide warmer at 60°C until fully dry. Slides were either used for downstream assays immediately after drying or stored at -80°C.

### Antigen retrieval of FFPE tissues, HOKs, and saliva for immunofluorescence, multiplexing, and RNAscope only

Slides with FFPE tissues, HOKs, and saliva were heated to 60°C on a slide warmer for 30 min, then immersed in a series of nonpolar to 100% nuclease-free water solutions (HistoChoice [Electron Microscopy Sciences] and 100% EtOH, 10 min; 90%, 70%, 50%, 30% EtOH in nuclease free water, 5 min each; 100% nuclease-free water, 10 min). For multiplexing, slides were then added to 1X citrate antigen retrieval buffer (pH 6; Sigma-Aldrich, C999, lot #MKCP6016) in nuclease-free water in a Coplin jar. Samples were antigen retrieved in a pressure cooker for 15 min at low pressure, then cooled to rt for at least 30 min before soaking in nuclease-free water (30 s), then 100% EtOH (3 min). For FFPE tissues and HOKs that were used for RNAscope only, these underwent the same protocol, apart from using 1X AR9 antigen retrieval buffer (pH 9; Akoya Biosciences).

### Immunofluorescence on HOKs and saliva

Blocking solution was prepared using gelatin from cold water fish skin (10%; Sigma-Aldrich #G7765-1L), normal donkey serum (10%; Jackson ImmunoResearch #017-000-121), bovine serum albumin (10%; Sigma-Aldrich A7030-50G), and Triton X-100 (0.2%; Sigma-Aldrich #T9284-500ML). After antigen retrieval (for gingival biopsies), a pap-pen was used to draw a hydrophobic barrier around the sample. Then, samples were washed for 2 × 5 min with RNAse-free 1X PBS before blocking using the blocking solution for 1h and incubated overnight at 4°C with antibody-doped blocking solution (Gt anti-LPS, polyclonal antibody RRID AB_1017872, ThermoFisher #PA1-73178, lot #ZF4350111, concentration; Ms anti-LTA, mAb 55, Hycult Biotech #HM5018-200UG, lot #37482M0124 1:400 concentration). After overnight incubation, the antibody-doped blocking solution was removed, and the sample was washed for 2 × 5 min with 1X PBS, then incubated with secondary antibody cocktail for 2h at rt (AlexaFluor 488 Dk anti-Gt, Jackson ImmunoResearch #705-545-003, lot #164823, 1:2000 concentration; Rhodamine Red-X Dk anti-Ms, Jackson ImmunoResearch #715-295-150, lot #156425, 1:500 concentration; Cyanine 5 Dk anti-Ms, Jackson ImmunoResearch #715-175-151, lot#155896, 1:400 concentration). The cocktail was then removed, and samples were washed with 1X PBS for 2 × 5 min, followed by a 5-min incubation with DAPI (1:1000 in 1X PBS) before washing with 1X PBS for 2 × 5 min and mounting with ProLong Gold Antifade (Invitrogen #P36930, Lot #2836845). Imaging of samples was performed using a Leica DMi8 with THUNDER Imager (Leica Microsystems) using a 20X NA 0.8 air or 40X or 63X 1.35 NA oil objectives.

### RNAscope HiPlex V2 on FFPE tissues and HOKs

All reagents in this section were purchased and used as received from ACD unless otherwise noted. HOKs were centrifuged using the MAPCELL methodology but did not undergo antigen retrieval prior to the protocol. The HybEZ oven was turned on and set to RNAscope (40°C), and the hydration paper was wetted with nuclease-free water. Following antigen retrieval, a hydrophobic barrier around the tissues was drawn using an ImmEdge pen (Vector Laboratories, #H-400, lot #ZG1113), and the slides were placed in the slide holder. One drop of Protease III reagent was added to each region containing sample; then, samples were placed in the HybEZ oven for 30 min. Slides were then washed in 1X RNAscope wash buffer (in nuclease-free water) before hybridizing with 1:100 of T probe solution for 2h (see Supplementary Data 2 for probe list). Samples were then subjected to a series of three amplification steps (30 min each with washing in-between steps) and FFPE reagent (2.5% reagent in 4X SSC; biopsies only) and fluorophore hybridization for 30 min each. After this series, RNAscope-provided DAPI was added to each sample for 30s, then tapped off the slides, followed by mounting using Prolong Gold Antifade (Invitrogen #P36930, Lot #2836845). Imaging of samples was performed using a Leica DMi8 with THUNDER Imager (Leica Microsystems) using a 40X or 63X 1.35 NA oil objective.

### Multiplexing of RNA and protein on the same slide

Primary and secondary antibodies were added to slides as described above. Following secondary antibody addition, samples were fixed with 10% NBF for 30 min in a covered humidity chamber. After 30 min, the NBF solution was pipetted off the slide, and the slide was washed with 1X PBS for 5 min. Following this washing step, the sample was used in the RNAscope assay. Imaging of multiplexed samples was performed using a Leica DMi8 with THUNDER Imager (Leica Microsystems) using a 40X or 63X 1.35 NA oil objective.

### Multi-scale imaging analysis of HOKs

We employed a dual staining approach with SYTO9 and DAPI to identify potential intracellular bacteria. HOKs immobilized using the MAPCELL methodology were stained with 0.1 μM SYTO9 and 0.3 μM DAPI in 1X PBS for 30 min. Subsequently, the samples were stained with 40 μg/mL wheat germ agglutinin (WGA) conjugated with Alexa Fluor 594 in 1× PBS for 10 min to label HGK cell plasma membranes. Post-staining, the slides were gently washed with 1× PBS once and mounted with ProLong Glass Antifade Mountant. The overview image was acquired with an Axiocam 705 mono camera with 10× 0.45 NA air objective (Zeiss), followed by super-resolution 3D confocal microscopy employing a LSM980 microscope with a 40× 1.2 NA water-immersion objective (Zeiss) with AiryScan 2. Sequential scanning was conducted using diode lasers (405, 488, and 561 nm), and the fluorescence emitted was collected with a AiryScan 2 detector*(60)*. AiryScan 3D deconvolution and multi-channel rendering (Max Intensity Projection) were performed using Zeiss Zen software and ImageJ FIJI, respectively.

## METAGENOMIC ANALYSES OF SINGLE-CELL RNA SEQUENCING DATA

### CellChat for analysis of incoming/outgoing and same-intracellular signaling strength

To quantify how cell-cell interactions between all cells shifted with bacterial elements, we used the R package *CellChat* (version 1.6.1, using the cell-cell interaction database). Previously, we created separate expression matrices for health and periodontitis from Tier 4 annotated AnnData objects*(20)*; from these, we created Seurat objects which we then input into *CellChat*. Analyses of these were conducted using CellChatDB, accounting for cell type composition differences. Bacterial state served as one of the variables in both health and periodontitis for which we compared to identify any significant changes.

### MultiNicheNet for bacterial association with keratinocytes: signatures and pathway enrichment

To understand how polybacterial intracellular macromolecules affected cell signaling pathways, we implemented MultiNicheNet (code available at https://github.com/saeyslab/multinichenetr) for analyses of our scRNAseq data. In a divergence from the main workflow, in both health and periodontitis, we considered bacterial state as a metavariable for analyses and filtered for KCs without bacterial state and KCs with bacterial state.

### Microbiome metagenomics of human oral keratinocytes (HOKs)

Raw paired-ends Fastq samples were preprocessed with Fastp*(61)* v0.22.0 with deduplication (-D) and low-complexity filtering (-y) and generating HTML and JSON reports before and after preprocessing. A first-pass removal of human reads aligning to the human genome reference GRCh38 was applied from Ensembl release 112 with Bowtie2*(62)* v2.5.3 with parameters “--very-sensitive-local”. Unmapped reads were collected back to Fastq files. Filtered Fastq reads were analyzed with the nf-core/taxprofiler:1.1.6 pipeline*(63)* with Nextflow v23.10.1 using Krakendb PlusPF, v20240112 for Kraken2*(64)* v2.1.2 and Bracken*(65)* v2.7 read classification and Krona for visualization (parameters “--run_kraken2 --run_brachen --run_krona”). Samples were analyzed separately to optimize compute resources.

Bracken output was used to compute Shannon’s diversity index with krakentools*(66)* v1.2 for each sample run. Indexes were combined and tested using a standard T-test. Taxonomic read counts were merged using custom scripts merging strains from the same species per sample and then normalized with edgeR*(67)* v4.0.16 using R v4.3.3. The expected dispersion was set to 0.16 (BCV=0.4) and exported normalized counts as a table for further analysis. When samples had contrasting categories, filtering was used for q-value < 0.05. Heatmaps were plotted using a log10(norm_counts) color scale without scaling.

### Keratinocyte analysis for predicted targets using Drug2Cell

To investigate the effects of drug treatment at single-cell resolution, we processed the single-cell RNA dataset of periodontium split by health and disease using the Scanpy library (v.1.98) in Python (v.3.9.12). The Drug2Cell (v0.1.0) framework was applied to predict drug-target interactions at the cellular level. Data were visualized using Matplotlib (v3.8.2) showing the mean expression of each cell type for a list of manually selected drug interactions.

## SUPPLEMENTAL FIGURE AND TABLE DESCRIPTIONS

**Figure 1S.**
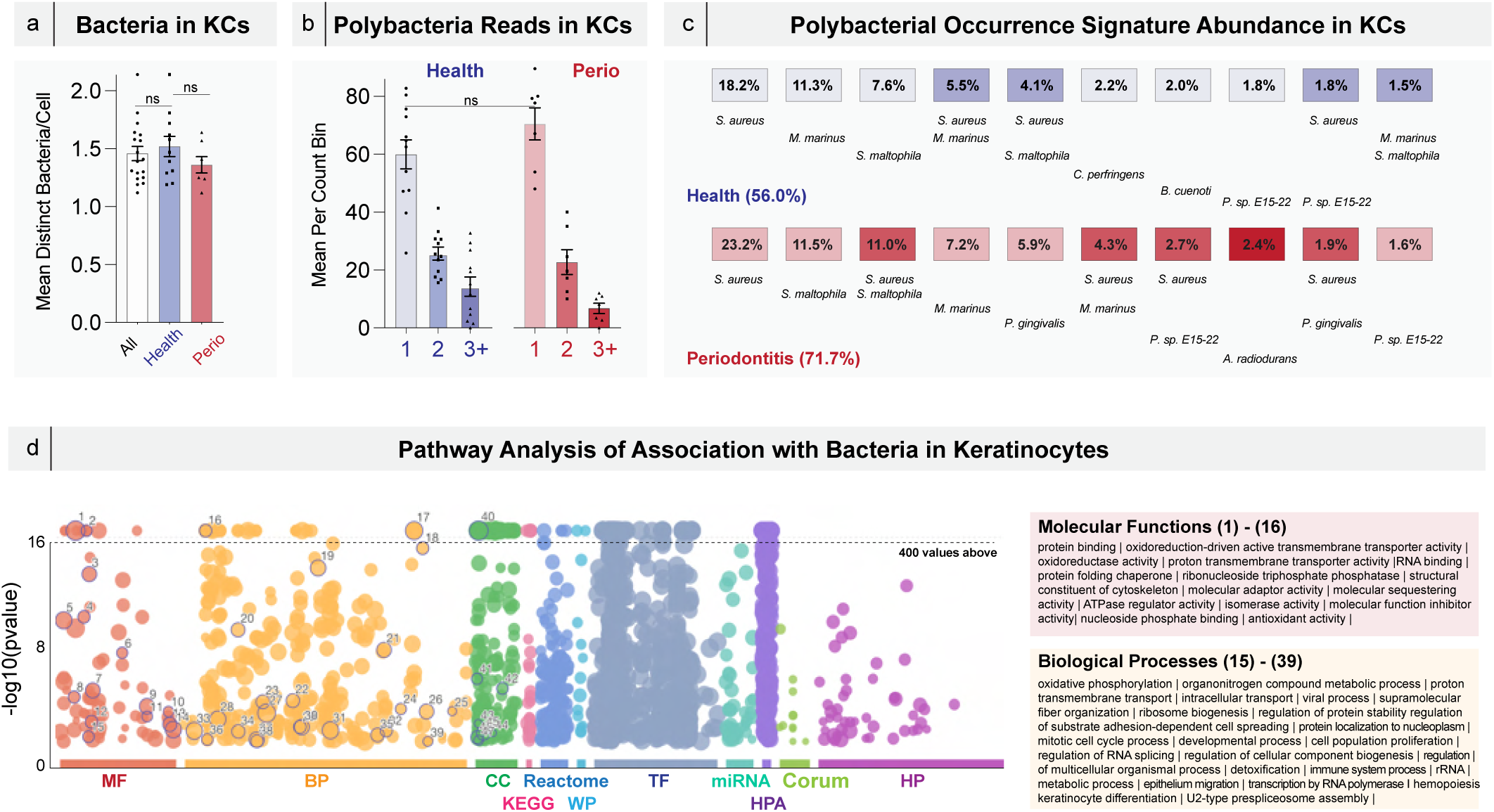
Bacteria counts, polybacterial counts, polybacterial occurrence signature abundance, and pathway analysis of association with bacteria in keratinocytes (KCs).

**Figure 2S.**
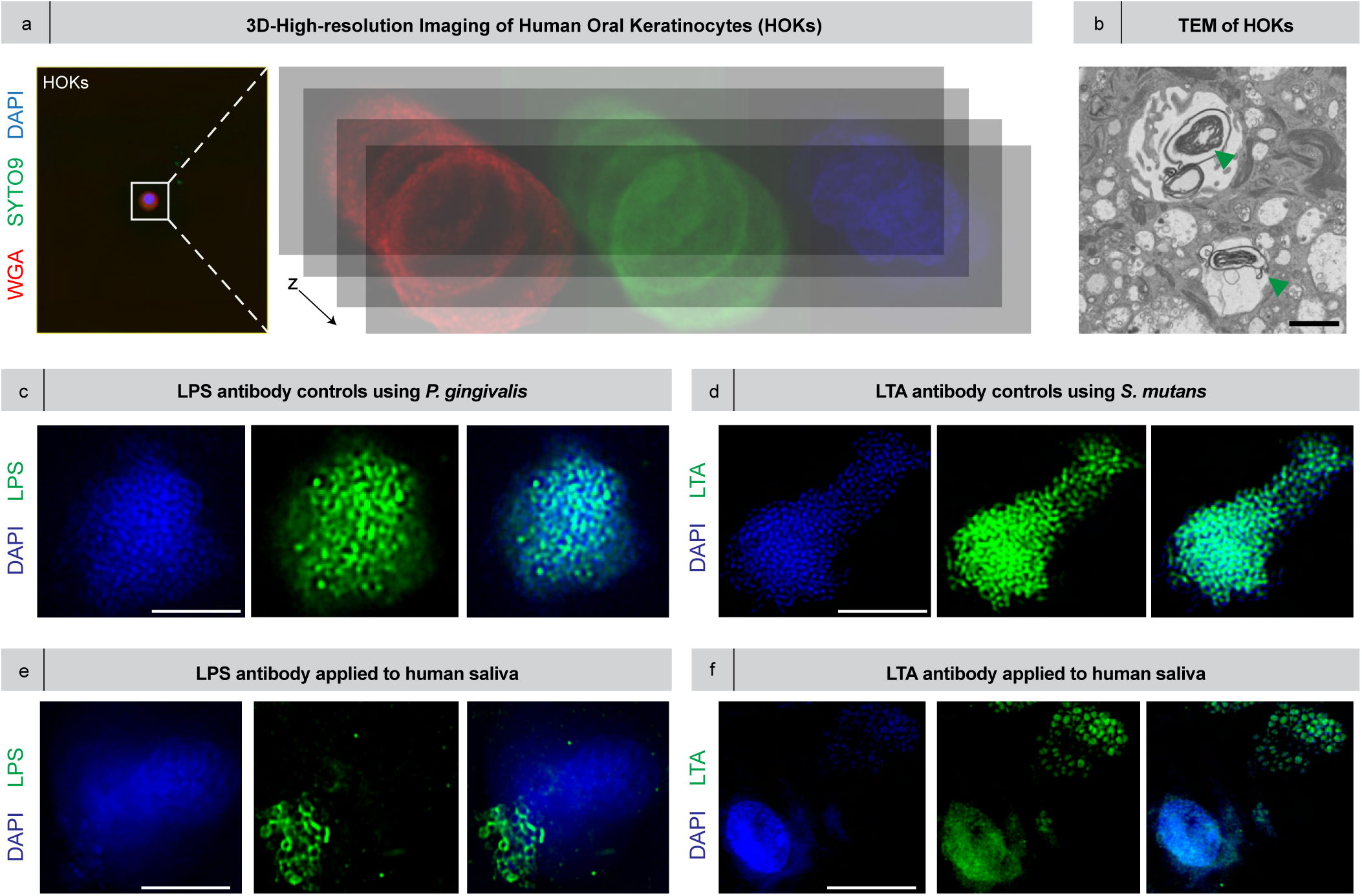
3D-High-resolution imaging of P4 human oral keratinocytes (HOKs), lipopolysaccharide (LPS) and lipoteichoic acid (LTA) antibody controls against pure bacterial cultures, and LPS and LTA antibodies applied to human saliva keratinocytes with naturally-associated bacteria.

**Figure 3S.**
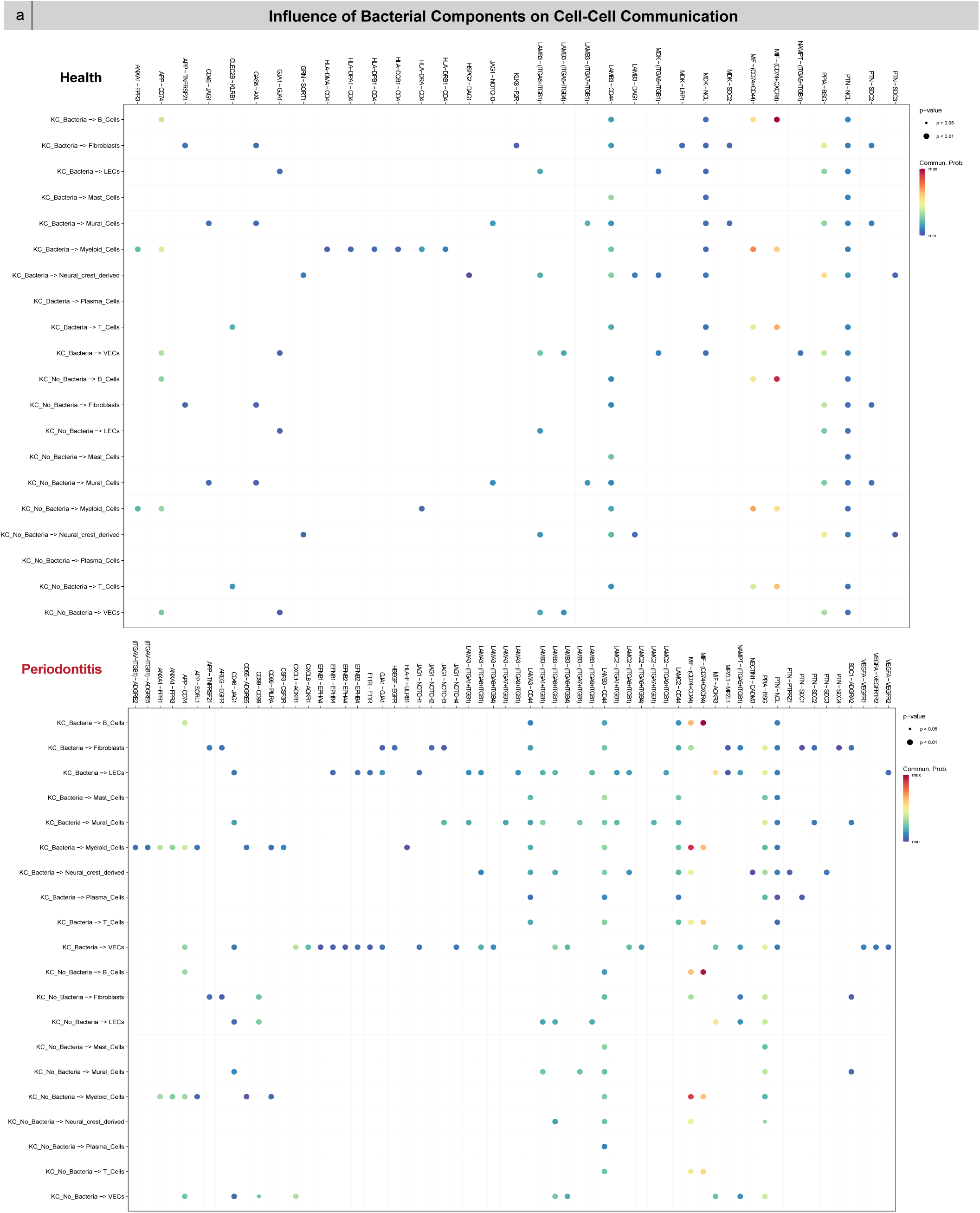
Influence of bacterial components on cell-cell communication.

**Figure 4S.**
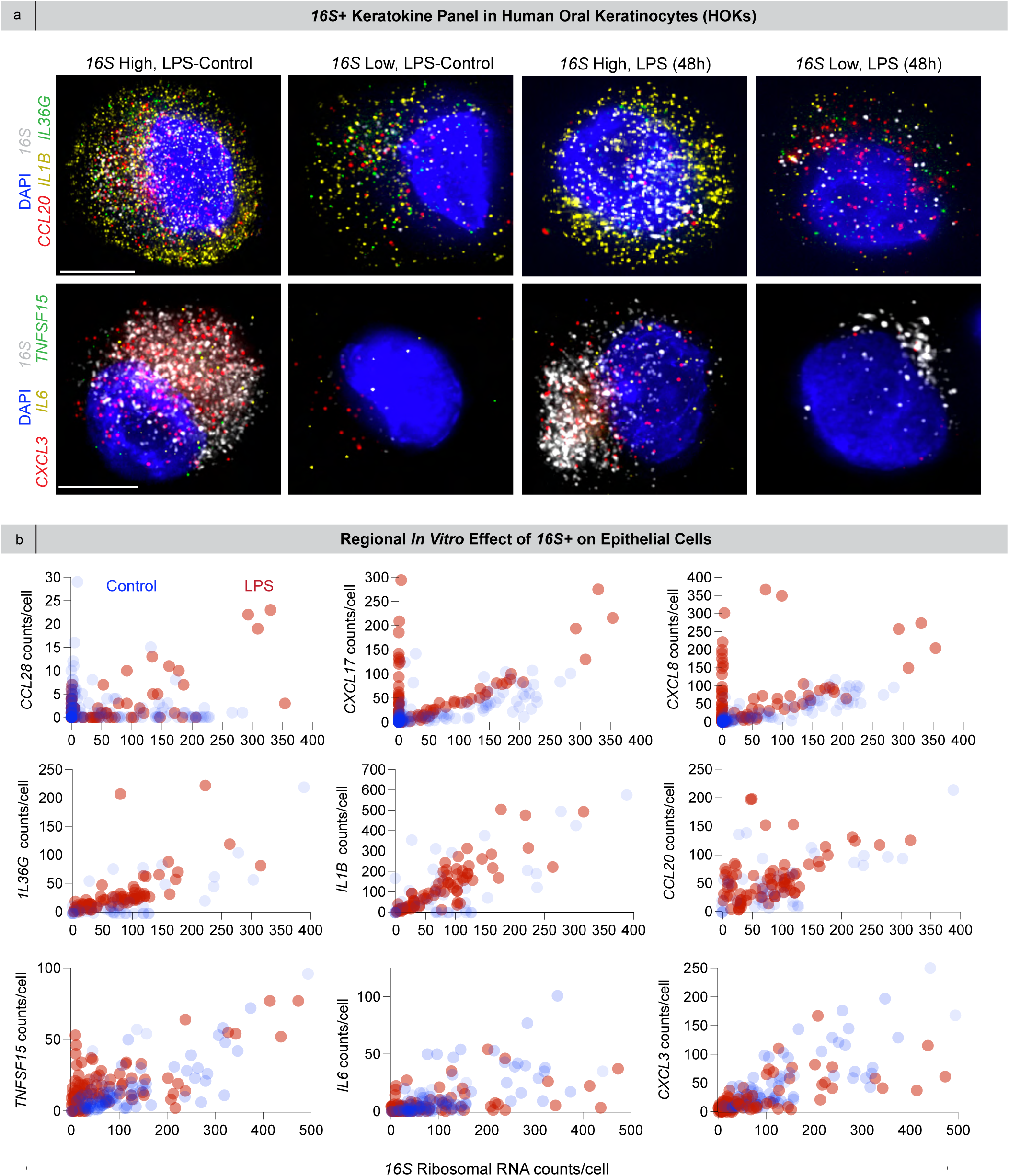
*16S*+ keratokine panel in HOKs and regional *in vitro* effect of *16S* on epithelial cells.

**Figure 5S.**
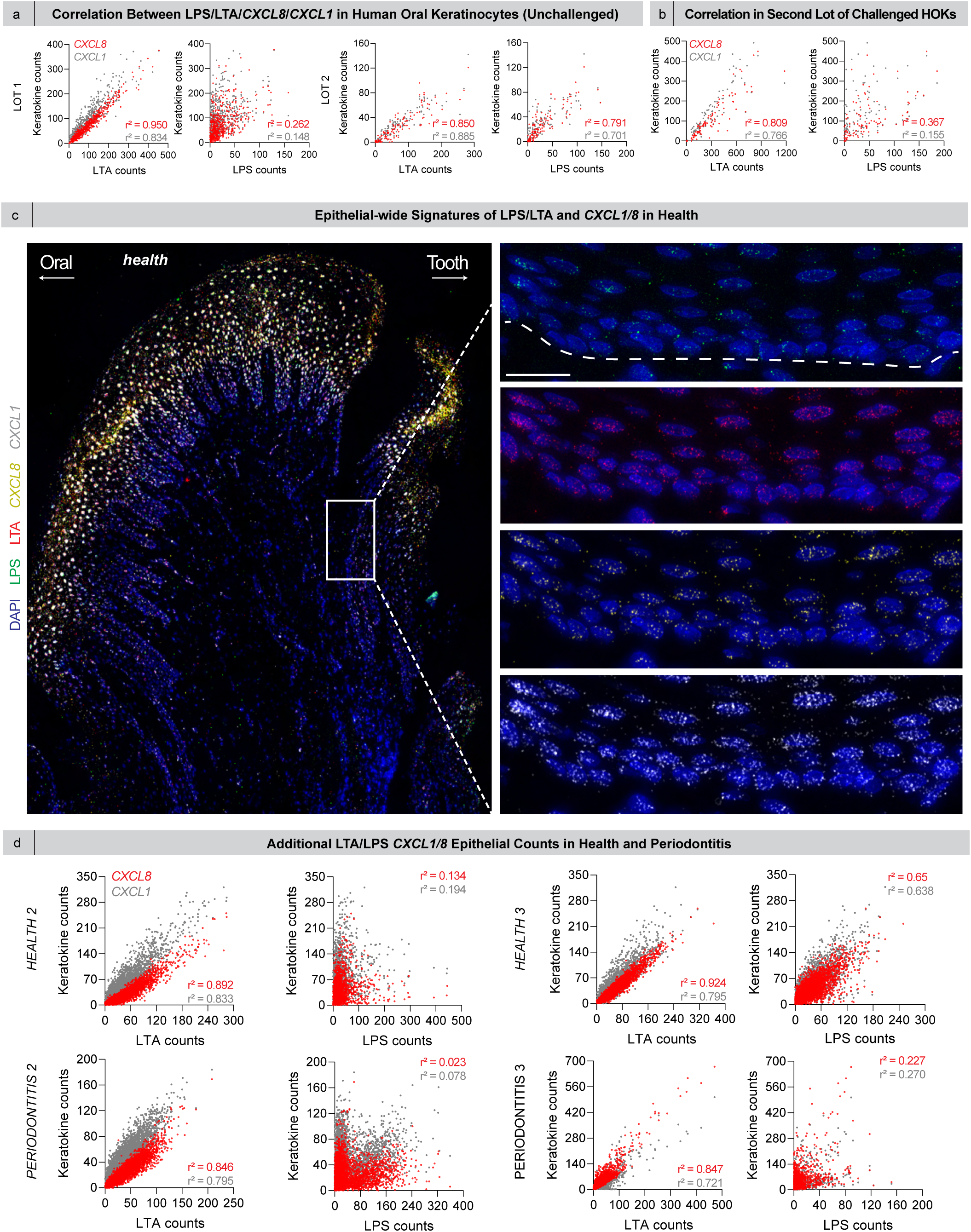
Correlation between LPS/LTA and *CXCL8/CXCL1* in unchallenged and challenged HOKs and in healthy and periodontitis gingival epithelia.

**Figure 6S.**
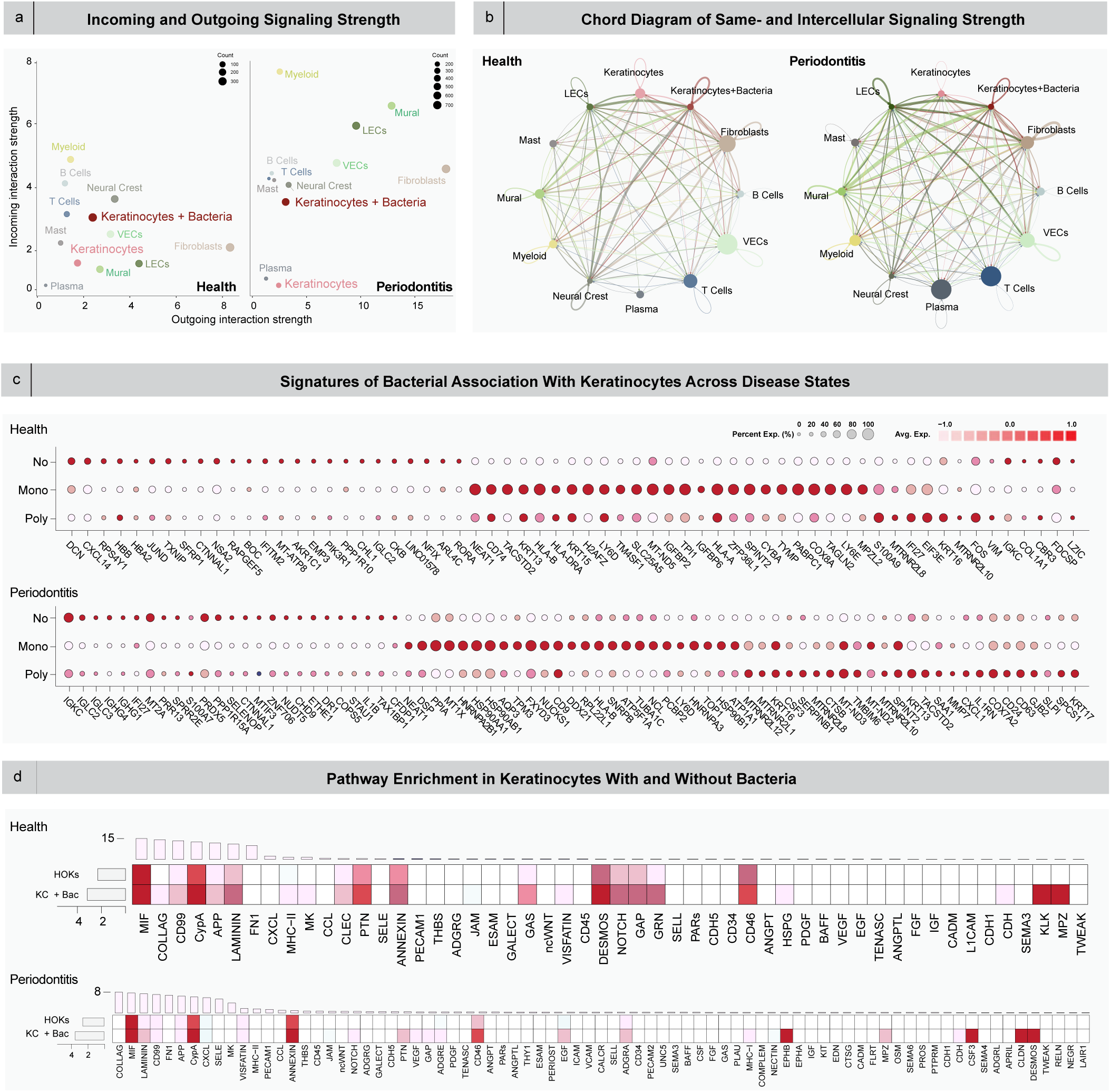
Incoming and outgoing signaling strength, same- and intracellular signaling strength, and bacterial association signatures with KCs and pathway enrichment in KCs across disease.

**Supplementary Data 1 |** Normalized and filtered counts of whole-genome sequencing results from HOKs and culture media.

**Supplementary Data 2 |** RNAscope probe list and healthy and periodontitis Drug2Cell ranks.

## References

1. N. M. Moutsopoulos, J. E. Konkel, Tissue-Specific Immunity at the Oral Mucosal Barrier. Trends Immunol 39, 276–287 (2018).

2. H. Kiyono, S. Fukuyama, NALT-versus Peyer’s-patch-mediated mucosal immunity. Nat Rev Immunol 4, 699–710 (2004).

3. I. F. Escapa, T. Chen, Y. Huang, P. Gajare, F. E. Dewhirst, K. P. Lemon, New Insights into Human Nostril Microbiome from the Expanded Human Oral Microbiome Database (eHOMD): a Resource for the Microbiome of the Human Aerodigestive Tract. mSystems 3 (2018), doi:10.1128/mSystems.00187-18.

4. F. E. Dewhirst, T. Chen, J. Izard, B. J. Paster, A. C. Tanner, W. H. Yu, A. Lakshmanan, W. G. Wade, The human oral microbiome. J Bacteriol 192, 5002–5017 (2010).

5. J. L. Baker, J. L. Mark Welch, K. M. Kauffman, J. S. McLean, X. He, The oral microbiome: diversity, biogeography and human health. Nat Rev Microbiol 22, 89–104 (2024).

6. X. Zhou, X. Shen, J. S. Johnson, D. J. Spakowicz, M. Agnello, W. Zhou, M. Avina, A. Honkala, F. Chleilat, S. J. Chen, K. Cha, S. Leopold, C. Zhu, L. Chen, L. Lyu, D. Hornburg, S. Wu, X. Zhang, C. Jiang, L. Jiang, L. Jiang, R. Jian, A. W. Brooks, M. Wang, K. Contrepois, P. Gao, S. M. S. Rose, T. D. B. Tran, H. Nguyen, A. Celli, B. Y. Hong, E. J. Bautista, Y. Dorsett, P. B. Kavathas, Y. Zhou, E. Sodergren, G. M. Weinstock, M. P. Snyder, Longitudinal profiling of the microbiome at four body sites reveals core stability and individualized dynamics during health and disease. Cell Host Microbe 32, 506–526 e9 (2024).

7. G. Hajishengallis, R. J. Lamont, H. Koo, Oral polymicrobial communities: Assembly, function, and impact on diseases. Cell Host Microbe 31, 528–538 (2023).

8. M. Hoggard, K. Biswas, M. Zoing, B. Wagner Mackenzie, M. W. Taylor, R. G. Douglas, Evidence of microbiota dysbiosis in chronic rhinosinusitis. Int Forum Allergy Rhinol 7, 230–239 (2017).

9. Y. J. Moon, J. H. Kim, J. W. Cho, J. Y. Na, T. I. Lee, D. Lee, D. Bae, E. Yoon, S. K. Kim, Microstructured void gratings for outcoupling deep-trap guided modes. Opt Express 26, A450–A461 (2018).

10. A. C. R. Tanner, C. A. Kressirer, S. Rothmiller, I. Johansson, N. I. Chalmers, The Caries Microbiome: Implications for Reversing Dysbiosis. Adv Dent Res 29, 78–85 (2018).

11. T. E. Van Dyke, P. M. Bartold, E. C. Reynolds, The Nexus Between Periodontal Inflammation and Dysbiosis. Front Immunol 11, 511 (2020).

12. Q. T. Easter, B. F. Matuck, B. M. Warner, K. M. Byrd, Biogeographical Impacts of Dental, Oral, and Craniofacial Microbial Reservoirs. J Dent Res 102, 1303–1314 (2023).

13. J. L. Mark Welch, F. E. Dewhirst, G. G. Borisy, Biogeography of the Oral Microbiome: The Site-Specialist Hypothesis. Annu Rev Microbiol 73, 335–358 (2019).

14. A. Acharya, Y. Chan, S. Kheur, L. J. Jin, R. M. Watt, N. Mattheos, Salivary microbiome in non-oral disease: A summary of evidence and commentary. Arch Oral Biol 83, 169–173 (2017).

15. D. Kim, J. P. Barraza, R. A. Arthur, A. Hara, K. Lewis, Y. Liu, E. L. Scisci, E. Hajishengallis, M. Whiteley, H. Koo, Spatial mapping of polymicrobial communities reveals a precise biogeography associated with human dental caries. Proc Natl Acad Sci U S A 117, 12375–12386 (2020).

16. M. Nagata, J. D. English, N. Ono, W. Ono, Diverse stem cells for periodontal tissue formation and regeneration. Genesis 60, e23495 (2022).

17. M. S. Tonetti, H. Greenwell, K. S. Kornman, Staging and grading of periodontitis: Framework and proposal of a new classification and case definition. J Periodontol 89 **Suppl 1**, S159–S172 (2018).

18. D. F. Kinane, P. G. Stathopoulou, P. N. Papapanou, Periodontal diseases. Nat Rev Dis Primers 3, 17038 (2017).

19. J. D. Beck, P. N. Papapanou, K. H. Philips, S. Offenbacher, Periodontal Medicine: 100 Years of Progress. J Dent Res 98, 1053–1062 (2019).

20. Q. T. Easter, B. Fernandes Matuck, G. Beldorati Stark, C. L. Worth, A. V Predeus, B. Fremin, K. Huynh, V. Ranganathan, Z. Ren, D. Pereira, B. T. Rupp, T. Weaver, K. Miller, P. Perez, A. Hasuike, Z. Chen, M. Bush, X. Qu, J. Lee, S. H. Randell, S. M. Wallet, I. Sequeira, H. Koo, K. M. Tyc, J. Liu, K. I. Ko, S. A. Teichmann, K. M. Byrd, Single-cell and spatially resolved interactomics of tooth-associated keratinocytes in periodontitis. Nat Commun 15, 5016 (2024).

21. A. J. Caetano, V. Yianni, A. Volponi, V. Booth, E. M. D’Agostino, P. Sharpe, Defining human mesenchymal and epithelial heterogeneity in response to oral inflammatory disease. Elife 10 (2021), doi:10.7554/eLife.62810.

22. D. W. Williams, T. Greenwell-Wild, L. Brenchley, N. Dutzan, A. Overmiller, A. P. Sawaya, S. Webb, D. Martin, N. N. Genomics, C. Computational Biology, G. Hajishengallis, K. Divaris, M. Morasso, M. Haniffa, N. M. Moutsopoulos, Human oral mucosa cell atlas reveals a stromal-neutrophil axis regulating tissue immunity. Cell 184, 4090–4104 e15 (2021).

23. N. Huang, P. Perez, T. Kato, Y. Mikami, K. Okuda, R. C. Gilmore, C. D. Conde, B. Gasmi, S. Stein, M. Beach, E. Pelayo, J. O. Maldonado, B. A. Lafont, S. I. Jang, N. Nasir, R. J. Padilla, V. A. Murrah, R. Maile, W. Lovell, S. M. Wallet, N. M. Bowman, S. L. Meinig, M. C. Wolfgang, S. N. Choudhury, M. Novotny, B. D. Aevermann, R. H. Scheuermann, G. Cannon, C. W. Anderson, R. E. Lee, J. T. Marchesan, M. Bush, M. Freire, A. J. Kimple, D. L. Herr, J. Rabin, A. Grazioli, S. Das, B. N. French, T. Pranzatelli, J. A. Chiorini, D. E. Kleiner, S. Pittaluga, S. M. Hewitt, P. D. Burbelo, D. Chertow, N. C.-A. Consortium, H. C. A. Oral, N. Craniofacial Biological, K. Frank, J. Lee, R. C. Boucher, S. A. Teichmann, B. M. Warner, K. M. Byrd, SARS-CoV-2 infection of the oral cavity and saliva. Nat Med 27, 892–903 (2021).

24. S. Ji, Y. Choi, Microbial and Host Factors That Affect Bacterial Invasion of the Gingiva. J Dent Res 99, 1013–1020 (2020).

25. J. A. Lemos, S. R. Palmer, L. Zeng, Z. T. Wen, J. K. Kajfasz, I. A. Freires, J. Abranches, L. J. Brady, The Biology of Streptococcus mutans. Microbiol Spectr 7 (2019), doi:10.1128/microbiolspec.GPP3-0051-2018.

26. S. Ji, J. E. Shin, Y. C. Kim, Y. Choi, Intracellular degradation of Fusobacterium nucleatum in human gingival epithelial cells. Mol Cells 30, 519–526 (2010).

27. T. Tominari, A. Sanada, R. Ichimaru, C. Matsumoto, M. Hirata, Y. Itoh, Y. Numabe, C. Miyaura, M. Inada, Gram-positive bacteria cell wall-derived lipoteichoic acid induces inflammatory alveolar bone loss through prostaglandin E production in osteoblasts. Sci Rep 11, 13353 (2021).

28. K. Kanemaru, J. Cranley, D. Muraro, A. M. A. Miranda, S. Y. Ho, A. Wilbrey-Clark, J. Patrick Pett, K. Polanski, L. Richardson, M. Litvinukova, N. Kumasaka, Y. Qin, Z. Jablonska, C. I. Semprich, L. Mach, M. Dabrowska, N. Richoz, L. Bolt, L. Mamanova, R. Kapuge, S. N. Barnett, S. Perera, C. Talavera-Lopez, I. Mulas, K. T. Mahbubani, L. Tuck, L. Wang, M. M. Huang, M. Prete, S. Pritchard, J. Dark, K. Saeb-Parsy, M. Patel, M. R. Clatworthy, N. Hubner, R. A. Chowdhury, M. Noseda, S. A. Teichmann, Spatially resolved multiomics of human cardiac niches. Nature 619, 801–810 (2023).

29. A. J. Caetano, O. Human Cell Atlas, B. Craniofacial, I. Sequeira, K. M. Byrd, A Roadmap for the Human Oral and Craniofacial Cell Atlas. J Dent Res 101, 1274–1288 (2022).

30. B. Ghaddar, M. J. Blaser, S. De, Denoising sparse microbial signals from single-cell sequencing of mammalian host tissues. Nat Comput Sci 3, 741–747 (2023).

31. F. E. Dewhirst, T. Chen, J. Izard, B. J. Paster, A. C. R. Tanner, W.-H. Yu, A. Lakshmanan, W. G. Wade, The human oral microbiome. J Bacteriol 192, 5002–17 (2010).

32. A. E. Burrows, A. Smogorzewska, S. J. Elledge, Polybromo-associated BRG1-associated factor components BRD7 and BAF180 are critical regulators of p53 required for induction of replicative senescence. Proc Natl Acad Sci U S A 107, 14280–14285 (2010).

33. J. Liang, L. Guo, K. Li, X. Xiao, W. Zhu, X. Zheng, J. Hu, H. Zhang, J. Cai, Y. Yu, Y. Tan, C. Li, X. Liu, C. Hu, Y. Liu, P. Qiu, X. Su, S. He, Y. Lin, G. Yan, Inhibition of the mevalonate pathway enhances cancer cell oncolysis mediated by M1 virus. Nat Commun 9, 1524 (2018).

34. S. Jin, C. F. Guerrero-Juarez, L. Zhang, I. Chang, R. Ramos, C. H. Kuan, P. Myung, M. V Plikus, Q. Nie, Inference and analysis of cell-cell communication using CellChat. Nat Commun 12, 1088 (2021).

35. S. T. Das, L. Rajagopalan, A. Guerrero-Plata, J. Sai, A. Richmond, R. P. Garofalo, K. Rajarathnam, Monomeric and dimeric CXCL8 are both essential for in vivo neutrophil recruitment. PLoS One 5, e11754 (2010).

36. E. Lowry, R. C. Chellappa, B. Penaranda, K. V Sawant, M. Wakamiya, R. P. Garofalo, K. Rajarathnam, CXCL17 is a proinflammatory chemokine and promotes neutrophil trafficking. J Leukoc Biol 115, 1177–1182 (2024).

37. C. Stringer, T. Wang, M. Michaelos, M. Pachitariu, Cellpose: a generalist algorithm for cellular segmentation. Nat Methods 18, 100–106 (2021).

38. Y. Zhan, R. Zhang, H. Lv, X. Song, X. Xu, L. Chai, W. Lv, Z. Shang, Y. Jiang, R. Zhang, Prioritization of candidate genes for periodontitis using multiple computational tools. J Periodontol 85, 1059–1069 (2014).

39. J. L. Galeano Nino, H. Wu, K. D. LaCourse, A. G. Kempchinsky, A. Baryiames, B. Barber, N. Futran, J. Houlton, C. Sather, E. Sicinska, A. Taylor, S. S. Minot, C. D. Johnston, S. Bullman, Effect of the intratumoral microbiota on spatial and cellular heterogeneity in cancer. Nature 611, 810–817 (2022).

40. J. L. Dzink, R. J. Gibbons, W. C. Childs 3rd, S. S. Socransky, The predominant cultivable microbiota of crevicular epithelial cells. Oral Microbiol Immunol 4, 1–5 (1989).

41. A. V Colombo, C. M. da Silva, A. Haffajee, A. P. Colombo, Identification of intracellular oral species within human crevicular epithelial cells from subjects with chronic periodontitis by fluorescence in situ hybridization. J Periodontal Res 42, 236–243 (2007).

42. P. Briaud, R. K. Carroll, Extracellular Vesicle Biogenesis and Functions in Gram-Positive Bacteria. Infect Immun 88 (2020), doi:10.1128/IAI.00433-20.

43. S. Chen, Q. Lei, X. Zou, D. Ma, The role and mechanisms of gram-negative bacterial outer membrane vesicles in inflammatory diseases. Front Immunol 14, 1157813 (2023).

44. M. G. Sartorio, E. J. Pardue, M. F. Feldman, M. F. Haurat, Bacterial Outer Membrane Vesicles: From Discovery to Applications. Annu Rev Microbiol 75, 609–630 (2021).

45. J. M. Bomberger, D. P. MacEachran, B. A. Coutermarsh, S. Ye, G. A. O’Toole, B. A. Stanton, Long-Distance Delivery of Bacterial Virulence Factors by Pseudomonas aeruginosa Outer Membrane Vesicles. PLoS Pathog 5, e1000382 (2009).

46. J. Jäger, S. Keese, M. Roessle, M. Steinert, A. B. Schromm, Fusion of Legionella pneumophila outer membrane vesicles with eukaryotic membrane systems is a mechanism to deliver pathogen factors to host cell membranes. Cell Microbiol 17, 607–20 (2015).

47. E. J. O’Donoghue, A. M. Krachler, Mechanisms of outer membrane vesicle entry into host cells. Cell Microbiol 18, 1508–1517 (2016).

48. G. Zhao, M. K. Jones, Role of Bacterial Extracellular Vesicles in Manipulating Infection. Infect Immun 91, e0043922 (2023).

49. K. Koeppen, T. H. Hampton, M. Jarek, M. Scharfe, S. A. Gerber, D. W. Mielcarz, E. G. Demers, E. L. Dolben, J. H. Hammond, D. A. Hogan, B. A. Stanton, A Novel Mechanism of Host-Pathogen Interaction through sRNA in Bacterial Outer Membrane Vesicles. PLoS Pathog 12, e1005672 (2016).

50. D. Li, M. Wu, Pattern recognition receptors in health and diseases. Signal Transduct Target Ther 6, 291 (2021).

51. J. Winter, D. Letley, J. Rhead, J. Atherton, K. Robinson, Helicobacter pylori membrane vesicles stimulate innate pro- and anti-inflammatory responses and induce apoptosis in Jurkat T cells. Infect Immun 82, 1372–1381 (2014).

52. P. Chandra, S. J. Grigsby, J. A. Philips, Immune evasion and provocation by Mycobacterium tuberculosis. Nat Rev Microbiol 20, 750–766 (2022).

53. E. G. Vozza, A. M. Kelly, C. M. Daly, S. A. O’Rourke, S. R. Carlile, B. Morris, A. Dunne, R. M. McLoughlin, Type 1 interferons promote Staphylococcus aureus nasal colonization by inducing phagocyte apoptosis. Cell Death Discov 10, 403 (2024).

54. J. Schindelin, I. Arganda-Carreras, E. Frise, V. Kaynig, M. Longair, T. Pietzsch, S. Preibisch, C. Rueden, S. Saalfeld, B. Schmid, J. Y. Tinevez, D. J. White, V. Hartenstein, K. Eliceiri, P. Tomancak, A. Cardona, Fiji: an open-source platform for biological-image analysis. Nat Methods 9, 676–682 (2012).

55. A. J. Marsh, M. A. Azcarate-Peril, M. R. Aljumaah, J. Neville, M. T. Perrin, L. L. Dean, M. D. Wheeler, I. N. Hines, R. Pawlak, Fatty acid profile driven by maternal diet is associated with the composition of human milk microbiota. Frontiers in Microbiomes 1 (2022), doi:10.3389/frmbi.2022.1041752.

56. A. A. Ribeiro, Y. Jiao, M. Girnary, T. Alves, L. Chen, A. Farrell, D. Wu, F. Teles, N. Inohara, K. V Swanson, J. T. Marchesan, Oral biofilm dysbiosis during experimental periodontitis. Mol Oral Microbiol 37, 256–265 (2022).

57. S. Sun, X. Zhu, X. Huang, H. J. Murff, R. M. Ness, D. L. Seidner, A. A. Sorgen, I. C. Blakley, C. Yu, Q. Dai, M. A. Azcarate-Peril, M. J. Shrubsole, A. A. Fodor, On the robustness of inference of association with the gut microbiota in stool, rectal swab and mucosal tissue samples. Sci Rep 11, 14828 (2021).

58. H. M. B. Seth-Smith, F. Bonfiglio, A. Cuénod, J. Reist, A. Egli, D. Wüthrich, Evaluation of Rapid Library Preparation Protocols for Whole Genome Sequencing Based Outbreak Investigation. Front Public Health 7, 241 (2019).

59. E. S. Reynolds, The use of lead citrate at high pH as an electron-opaque stain in electron microscopy. J Cell Biol 17, 208–12 (1963).

60. X. Wu, J. A. Hammer, ZEISS Airyscan: Optimizing Usage for Fast, Gentle, Super-Resolution Imaging. Methods Mol Biol 2304, 111–130 (2021).

61. S. Chen, Ultrafast one-pass FASTQ data preprocessing, quality control, and deduplication using fastp. iMeta 2, e107 (2023).

62. B. Langmead, S. L. Salzberg, Fast gapped-read alignment with Bowtie 2. Nat Methods 9, 357–9 (2012).

63. S. Stamouli, M. E. Beber, T. Normark, L. Andersson-Li, M. Borry, M. Jamy, J. A. Fellows Yates, nf-core community. bioRxiv, 1–28 (2023).

64. D. E. Wood, J. Lu, B. Langmead, Improved metagenomic analysis with Kraken 2. Genome Biol 20, 257 (2019).

65. J. Lu, F. P. Breitwieser, P. Thielen, S. L. Salzberg, Bracken: Estimating species abundance in metagenomics data. PeerJ Comput Sci 2017 (2017), doi:10.7717/peerj-cs.104.

66. J. Lu, N. Rincon, D. E. Wood, F. P. Breitwieser, C. Pockrandt, B. Langmead, S. L. Salzberg, M. Steinegger, Author Correction: Metagenome analysis using the Kraken software suite. Nat Protoc (2024), doi:10.1038/s41596-024-01064-1.

67. Y. Chen, A. T. L. Lun, G. K. Smyth, From reads to genes to pathways: differential expression analysis of RNA-Seq experiments using Rsubread and the edgeR quasi-likelihood pipeline. F1000Res 5, 1438 (2016).

